# USP15 REGULATES NEUROINFLAMMATION AND DRIVES PATHOGENESIS IN SYNUCLEINOPATHIES

**DOI:** 10.64898/2026.09.05.749595

**Authors:** Nassima Fodil, Esther Del Cid-Pellitero, Valerio E.C. Piscopo, Julia Xiao Xuan Luo, Michael Wong, Jean Frederic Olivier, Aeshah Alluli, Irina Shlaifer, Zhipeng You, Carol X-Q Chen, Nathalia Aprahamian, Yoonjeong Cha, Étienne Collette, Jonas J Mayo, Alva Annett, Nada Jabado, Claudia L Kleinman, Mohamedi N. Kagalwala, Jermaine Ross, Mark Lathrop, David Langlais, Thomas M. Durcan, Edward A. Fon, Philippe Gros

## Abstract

Neuroinflammation strongly contributes to the pathogenesis of neurological and neurodegenerative diseases, including Parkinson’s disease (PD). We show that ablation of *Usp15* in astrocytes and in microglia protects against lethal neuroinflammation in vivo. In a mouse model of synucleinopathy, *Usp15* deletion diminishes α-syn deposits in the brain, slows disease progression, and increases survival time. The neuroprotective effect of *Usp15* is associated with differential expression of inflammatory pathways *in situ* including interferon stimulated genes. These USP15-dependent effects *in vivo* are recapitulated *in vitro* in primary human microglia and astrocytes. We detect high USP15 expression in microglia from Parkinson’s patients with strong co-expression with LRRK2 and SNCA. In humans, we detect a strong cis-acting eQTL directing high USP15 expression in CD14+ myeloid cells. The allele driving this eQTL is itself associated with increased disease risk, linking myeloid USP15 expression, elevated USP15 plasma levels in Parkinson’s patients, to genetic susceptibility.

## INTRODUCTION

Parkinson’s Disease (PD) is a neurodegenerative disorder characterized by the appearance of intracellular aggregates (Lewy bodies) of fibrillar forms of α-synuclein (α-syn) that accumulate in neuronal bodies and neurites, leading to progressive degeneration and loss of dopaminergic neurons in the midbrain substantia nigra, and appearance of motor symptoms, including bradykinesia, resting tremor, rigidity, and postural instability^1, 2^. Several factors have been implicated in PD pathogenesis at the cellular and molecular levels, including mitochondrial dysfunction^3, 4^, oxidative stress^5, 6^, environmental exposure ^7, 8^, pre-disposing genetic factors (variants in SNCA (α-syn), LRRK2, PRKN, PINK1, DJ-1, and VPS35^3, 9, 10, 11, 12, 13^) and neuroinflammation (NI)^14, 15, 16^. Genome-wide association studies (GWAS) have mapped several genetic variants associated with PD risk, many of which are located at or near immune-related genes^17, 18, 19^, supporting a role for NI in onset and progression of PD. Such pathological NI is characterized by elevated levels of pro-inflammatory cytokines and chemokines in the central nervous system (CNS)^20^, glial cell activation *in situ*^21^, with further exacerbation by infiltrating activated myeloid and lymphoid cells^22^. Studies of post-mortem brains from PD patients have indeed documented the presence of activated microglia (up-regulation of HLA-DR, CD68, and TLRs), suggesting a key contribution of these cells to NI^23^.

Microglia and astrocytes are glial cells whose normal function is essential to maintain CNS homeostasis^24^. Microglia can become chronically activated in response to persistent cell damage and other stimuli, which causes NI and drives PD pathogenesis^25, 26, 27^. For instance, α-syn aggregates lead to the release of pro-inflammatory cytokines (e.g., TNF-α, IL-1β, IL-6), reactive oxygen species, and nitric oxide, which further exacerbate neuronal damage^26^. Furthermore, inhibition of microglial reactivity in mouse models of PD delays the progression of α-syn pathology^28^. Similarly, in PD, astrocytes undergo a phenotypic shift toward a reactive state^29^. Reactive astrocytes can release inflammatory mediators and transition to a neurotoxic A1 phenotype, contributing to neuronal loss and disease progression^30^. Moreover, crosstalk between microglia and astrocytes can amplify the inflammatory milieu, fostering a toxic environment for neurons ^30^. Indeed, the inhibition of microglial-mediated conversion of astrocytes to an A1 neurotoxic phenotype prevents α-synucleinopathy in a mouse model of PD^31^.

Genetic studies in mouse models have identified *Usp15* (de-ubiquitinase family member) as a critical regulator of NI ^32^. Mutational inactivation of *Usp15* (*Usp15^L749R^* or *Usp15^KO/KO^*) protects against lethal experimental cerebral malaria (ECM) and experimental autoimmune encephalomyelitis (EAE)^33^. *Usp15* was found to be expressed in both CNS-resident cells (microglia and astrocytes) and infiltrating immune cells, underscoring its broad relevance to neuroinflammatory signalling networks^33^.

*Usp15* appears most critical for activation of RIG-I dependent type I interferon (Type I IFN) signaling pathways which includes expression of Interferon Stimulated Genes (ISG) in response to viral dsRNA and other PAMPs^32, 34, 35^. The RIG-I signaling complex comprises 2 ligand binding CARD domains whose tetramerization upon signaling induces recruitment of the MAVS adaptor. The oligomerized CARD tetramers are stabilized by ubiquitination via the E3 ubiquitin ligase TRIM25 to enhance MAVS engagement, stimulation of the kinases TBK1 and IKKε, phosphorylation of IRF3 and IRF7, and transcription of Type I IFNs and associated ISGs^34, 35^. Using biochemical studies, we showed that USP15 binds to and de-ubiquitinates TRIM25 ^33^. Furthermore, inactivation of *Trim25* in mice (*Trim25^KO/KO^*) blunts NI in the ECM model; likewise, *Trim25^KO/+^:Usp15^L749R/+^* compound heterozygotes are also resistant to NI, showing genetic complementation between *Usp15* and *Trim25*. Furthermore, we showed that *Usp15* and *Trim25* are co-expressed in mouse primary myeloid cells, and that their genetic inactivation dampens ISG gene expression in response to poly(I:C) and 3pRNA. Finally, USP15 and TRIM25 are also co-expressed in primary human microglia and astrocytes where they are up-regulated by IFN-γ, TNF-α and IL-1β^33^. Together, these findings show that TRIM25 and USP15 genetically, physically and functionally interact to positively regulate RIG-I dependent Type I IFN responses.

Here, we used a combination of in vivo studies in mouse models of NI and neurodegeneration, transcriptomics profiling by bulk RNA-seq and studies in human iPSC-derived microglia and astrocytes to identify and characterize the role of USP15 in the onset and progression of PD. We identify the cell types and molecular pathways involved, which together strongly support a role for USP15-driven activation of inflammatory pathways during the pathogenesis of PD. In support of *USP15* as a candidate PD gene, the human chr12q14.1 locus that contains *USP15* shows suggestive but sub-genome-wide-significant association in the largest European PD meta-GWAS to date (lead variant rs4763185, *P* = 3.0×10⁻⁶). *USP15* also receives a positive Polygenic Priority Score (PoPS = +0.256) in a recent genome-wide gene-prioritization analysis^36^,. Here, we extend this human-genetic evidence with cis-eQTL colocalization, conditional fine-mapping and Mendelian randomization, providing increased evidence supporting a role of the locus in susceptibility to PD.

## RESULTS

### *Usp15* Drives Neuroinflammation via Brain-Resident Immune Cells

To investigate the cell type responsible for the *Usp15*-regulated NI in the CNS, we generated a mouse bearing a conditional null allele at *Usp15* (*Usp15 ^flox/flox^*), which was crossed to transgenic lines carrying a tamoxifen-inducible Cre recombinase under the control of astrocytes (*Aldh1l1*-Cre) and microglia/monocyte/macrophage (*Cx3cr1*-Cre) specific promoters (Figure 1A). Following Cre induction, mice were infected with *Plasmodium berghei* ANKA (PbA) to induce cerebral malaria-associated lethal NI (Figure 1B). Results show that both astrocyte-specific and microglia-specific deletion of *Usp15* reduced clinical signs of NI and significantly improved (p<0.0001) survival compared to non-excised control littermates (Figure 1C).

**Figure 1.**
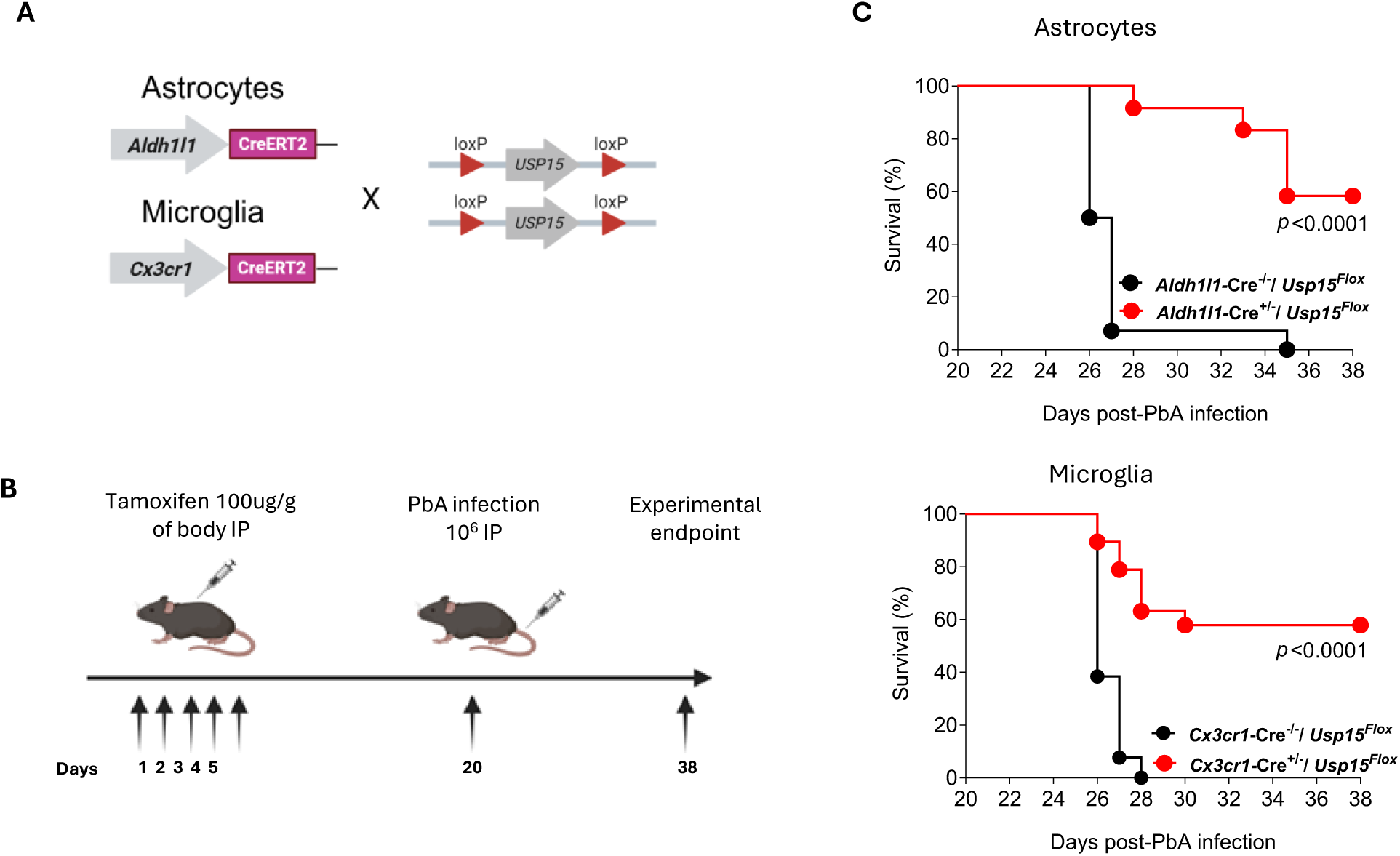
USP15 deficiency in microglia or astrocytes protects mice against lethal cerebral malaria in *Plasmodium berghei* ANKA infection. **(A)** Schematic representation of the strategy for cell-specific, Cre-mediated inactivation of USP15. *Usp15^flox/flox^* mice were crossed to *Aldh1l1*-CreERT2 mice to achieve astrocyte-specific deletion or to *Cx3cr1*-CreERT2 mice to achieve microglia-specific deletion following tamoxifen induction. **(B)** Experimental timeline. Mice received tamoxifen (100 µg/g body weight, intraperitoneally, 5 consecutive days) to induce Cre-mediated recombination. Two weeks after the last tamoxifen treatment, mice were infected intravenously with 10^6^ *Plasmodium berghei ANKA* parasitized red blood cells (Pba infection), and survival was monitored up to day 18 post-infection (corresponding to day 38 post tamoxifen treatment). **(C)** Kaplan–Meier plots depicting survival following PbA infection. Astrocyte-specific (*Aldh1l1*-Cre × *Usp15^flox/flox^*) and microglia-specific deletion of *Usp15* (*Cx3cr1*-Cre × *Usp15^flox/flox^*) significantly improved survival compared with Cre-negative littermate controls (p < 0.0001).

To examine the cellular and molecular pathways regulated by *Usp15* in astrocytes and microglia during acute NI *in vivo*, we conducted bulk RNA sequencing (RNA-seq) from brains of mice deleted for *Usp15* expression in astrocytes (*Aldh1l1*-Cre) or in microglia (*Cx3cr1*-Cre), 5 days after infection with PbA, and examined the genes differentially expressed (DEG) compared to WT controls (Figure 2A). Principal component analysis (PCA) identified two components contributing to variability in the dataset. The strongest effect was caused by infection (PC1), with a smaller effect contributed by the USP15 genotype (PC2). This genotype PC2 component was more evident in brains of animals with conditional deletion of *Usp15* in astrocytes compared to microglia (Figure 2B). Preliminary analysis of DEGs (GO term, Figure 2C) of the WT brains show the expected inflammatory response following PbA infection (Figure 2C). This response was attenuated in *Usp15* conditionally ablated mice, especially in astrocytes (*Aldh1l1-Cre*) that showed marked attenuation of multiple hallmark inflammatory pathways in the CNS (Figure 2C), including interferon programs, allograft rejection signatures, the IL-6–JAK–STAT3 axis, TNF-α signaling via NF-κB, apoptotic pathways, and complement activation. Differential expression of interferon-stimulated genes (ISGs), including *Ifit3*, *Irf7*, *Mx1*, *Oas2*, and *Stat1* was particularly evident (Supplemental Figure 1). Gene set enrichment analysis (GSEA) further showed that both type I and II interferon pathways were significantly reduced in both astrocytes (*Aldh1l1-Cre*) and microglia (*Cx3cr1*-Cre) *Usp15* deleted mice when compared to WT PbA-infected brains (Figure 2D).

**Figure 2.**
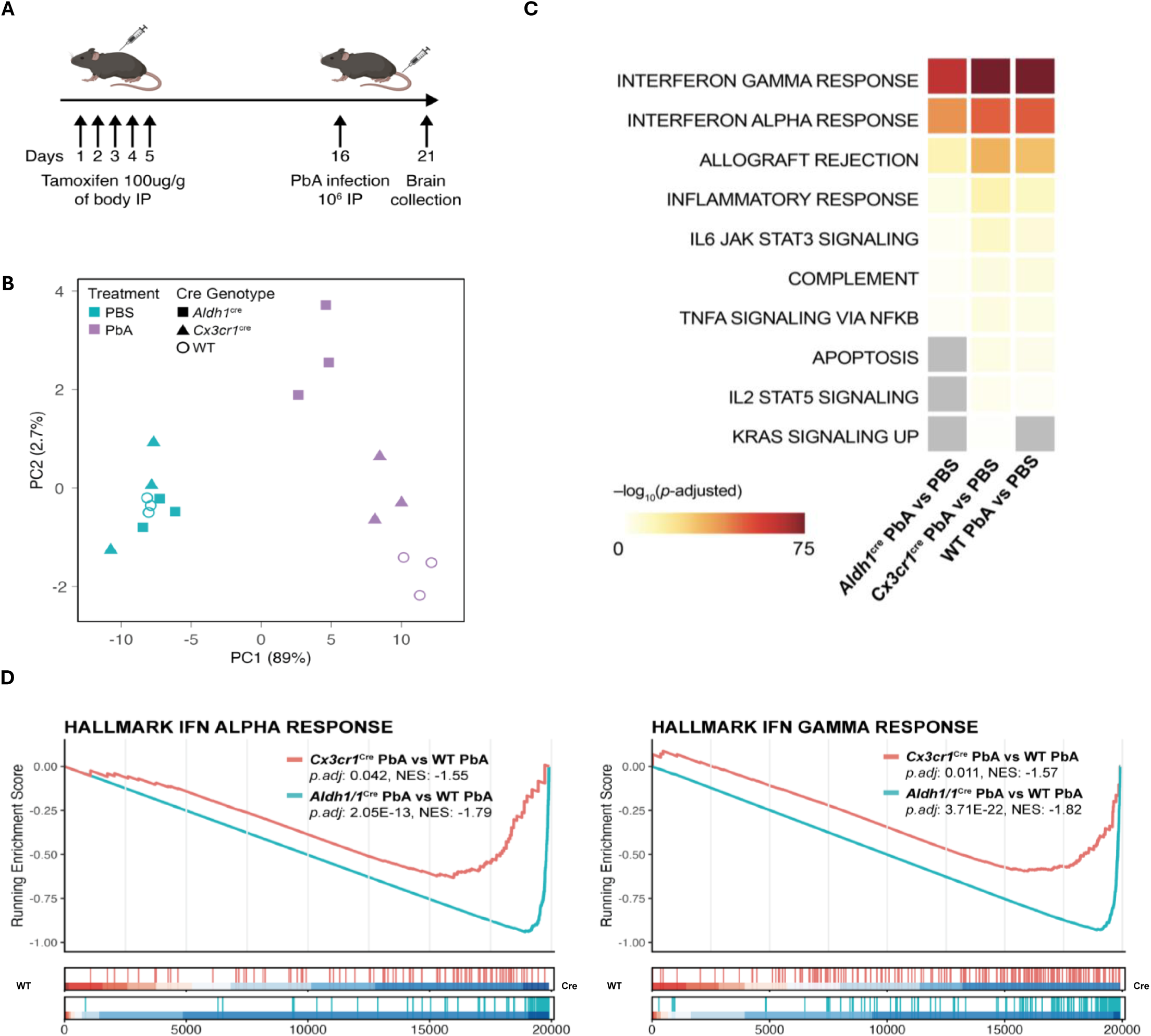
Bulk RNA-seq analysis of *Plasmodium berghei*–infected brains reveals cell-specific effects of USP15 deficiency in astrocytes and microglia. **(A)** Experimental design for bulk RNA-seq. Mice conditionally inactivated for Usp15 in astrocytes (*Aldh1l1*-CreERT2) and microglia (*Cx3cr1*-CreERT2), along with wild-type littermate controls, were treated with tamoxifen for five consecutive days followed by PbA infection on day 16, and brains were collected at day 5 post-infection (day 21 from start of tamoxifen treatment). **(B)** Principal component analysis (PCA) of bulk brain RNA-seq data. Samples segregated strongly by infection status along PC1, with additional genotype-dependent separation within the PbA-infected group. **(C)** Gene set enrichment analysis (GSEA) of differentially expressed genes comparing PbA-infected versus PBS-treated brains across genotypes. Hallmark inflammatory pathways, including type I and type II interferon responses, allograft rejection, IL-6–JAK–STAT3 signaling, TNFα signaling via NF-κB, apoptosis, and complement activation were induced in infected WT brains but attenuated in *Usp15*-deficient mice. Heatmap is colored by −log₁₀(adjusted p value); grey indicates non-significant enrichment. **(D)** GSEA running enrichment scores graphs for interferon alpha and interferon gamma response pathways showing decreased enrichment in USP15-deficient mice.

Together, these results show that inactivation of *Usp15* in astrocytes or microglia dampens expression of neuroinflammatory transcriptional programs in the brain (cytokines, chemokines, and ISGs) during acute lethal NI, blunting pathogenesis and increasing survival.

### *Usp15* Inhibition Reduces Neuroinflammation and Improves Outcomes in a Mouse Model of PD

We further examined a possible contribution of *Usp15* to the establishment of chronic states of NI driving pathogenesis in neurodegenerative diseases. For this, we used the M83 mouse strain, a model of α-synucleinopathy and PD^37^. M83 transgenic mice carry multiple copies of the pathogenic mutant variant of the human α-syn gene (hα-syn-A53T; from familial cases of PD) that enhances α-syn misfolding and aggregation, producing Lewy body-like pathology. Intra-striatal injection of preformed fibrils of α-syn (PFFs) in M83 mice causes an accelerated pathology which recapitulates key features of synucleinopathy and PD^38^.

Hemizygous M83 mice carrying the loss-of-function *Usp15* mutation (*Usp15^L749R^*; *Usp15^mut^*) were generated and, together with wild-type littermates (*Usp15^+/+^*), were injected with PFF, and the effect on pathogenesis was assessed by time of appearance and duration of neurological symptoms (motor impairment, body weight loss), and overall survival (Figure 3). Inactivation of *Usp15* significantly slowed disease progression (Figure 3B; number of days with clinical symptoms; *p* < 0.0001, Mann-Whitney test),and significantly increased overall survival (Figure 3C; *p* < 0.0001, Log-rank test), indicating a significant attenuation of neurodegeneration progression. Consistent with improved survival and clinical scores, *Usp15* mutant mice showed reduced accumulation of phospho-synuclein (pSyn) aggregates in the brain at day 84 in the substantia nigra and pons (Figure 3D, E). *Usp15*-mutant mice displayed fewer and less compact pSyn inclusions in the substantia nigra (Figure 3D) and reduced aggregation in the pons (Figure 3E). Collectively, these findings suggest that *Usp15* promotes α-syn propagation and NI, contributing to synucleinopathy progression *in vivo*.

**Figure 3.**
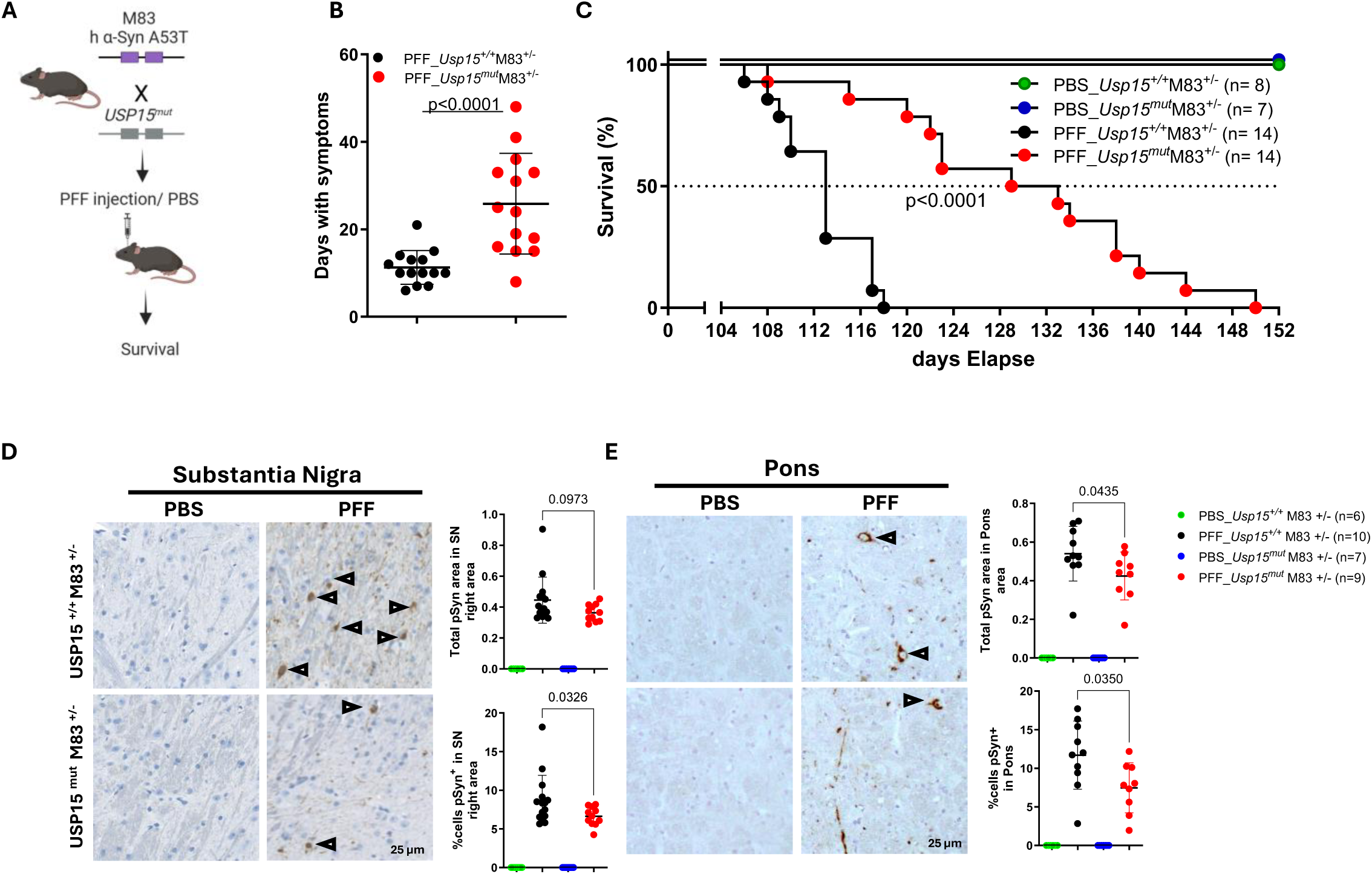
Ablation of USP15 protects mice against α-syn–induced neurodegeneration in the M83 Parkinson’s disease model. **(A**) Breeding scheme and experimental design. M83^+/−^transgenic mice expressing human A53T α-syn (hα-syn-A53T) were crossed with mice carrying a loss-of-function *Usp15* L749R mutation (USP15^mut^). Adult *Usp15^mut^*M83 mice and wild-type (*Usp15^+/+^*) littermates received intrastriatal injections of α-syn preformed fibrils (PFFs) or PBS and were monitored longitudinally for onset of symptoms and survival. **(B)** Quantification of disease progression. *Usp15^mut^*M83 mice displayed a significant delay in appearance of symptoms and duration of symptomatic period compared to *Usp15^+/+^*M83 littermates (Mann– Whitney test, p < 0.0001), indicating attenuation of neurodegenerative progression. PBS-injected mice of any genotype did not develop symptoms. **(C)** Kaplan–Meier survival analysis following intrastriatal PFF injection. *Usp15^mut^*M83 mice exhibited significantly prolonged survival compared with *Usp15^+/+^*M83 controls (Log-rank Mantel–Cox test, p < 0.0001). **(D)** Immunohistochemical analysis of phosphorylated α-syn (pSer129-α-syn; pSyn) accumulation in the substantia nigra (SN) 84 days after PFF injection. Representative images and quantification show extensive pSyn-positive neuronal inclusions in PFF-treated *Usp15^+/+^*M83 mice, whereas *Usp15^mut^*M83 mice exhibit a reduced pSyn burden, reflected by decreased total pSyn-positive area (top quantification) and a lower percentage of pSyn-positive neurons (bottom quantification) **(E)** Same as panel **(D)** but for the Pons area of the brain. Scale bars, 25 µm.

### Loss of *Usp15* Dampens Neuroinflammatory Interferon Signaling during α-synucleinopathy

We used bulk RNA-seq to identify molecular pathways regulated in the brain by *Usp15* and determine whether their dampening may underlie neuroprotection against neurodegeneration in M83 mice. These would be detected as differentially expressed genes (*Usp15^+/+^* versus *Usp15^mut^*) during α-synucleinopathy pathogenesis at an early point (84 days post PFF treatment), preceding the appearance of clinical symptoms in either group (>100d) (Figure 4A). PCA revealed that the two most contributing factors in the gene expression variance were PFF treatment (PC1) and genotype (PC2), which contributed ∼16% and ∼11% of variability in the dataset, respectively (Figure 4B). PFF treatment induced a larger transcriptional response in WT mice (*Usp15*^+/+^, 145 upregulated genes) compared to mutants (*Usp15^mut^*, 98 upregulated genes) (Supplemental Figure 2A). Examination of DEGs (cut-off of 1.5 fold; adjusted P value <0.05) identified 74 genes that were induced by treatment in both genotypes, while 73 genes were preferentially induced in *Usp15*^+/+^ mice compared to *Usp15^mut^* mutants (Supplemental Figure 2B-C).

**Figure 4.**
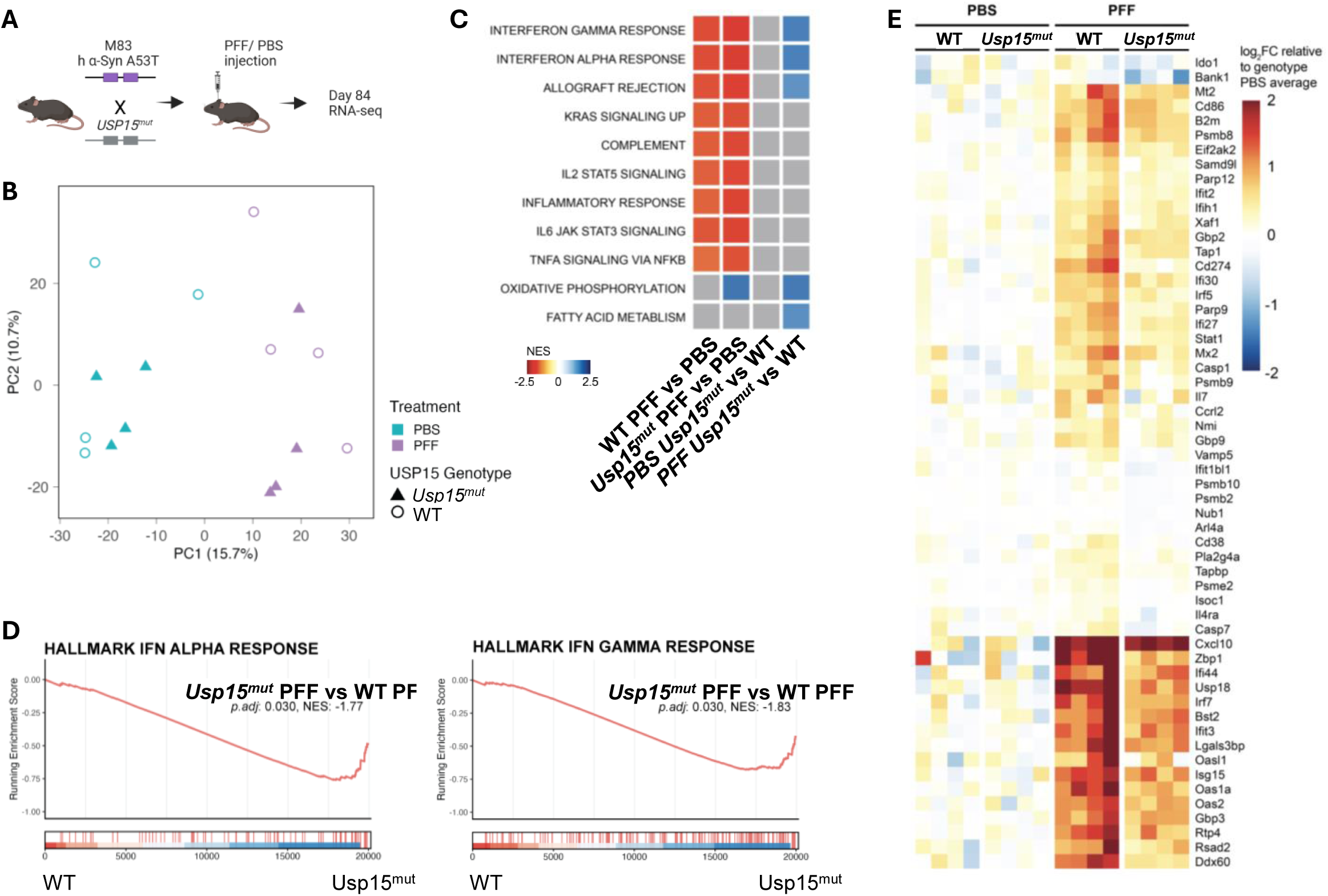
USP15 is required for interferon-driven neuroinflammatory transcriptional responses to α-syn pathology. **(A)** Experimental design. M83 transgenic mice expressing human A53T α-syn (hα-syn-A53T) carrying either a loss-of-function *Usp15* mutation (*Usp15^mut^*) or wild-type alleles were subjected to intrastriatal injection of α-syn preformed fibrils (PFFs) or PBS. Whole brains were collected 84 days post-injection for bulk RNA-seq. **(B)** Principal component analysis (PCA) of brain transcriptomes. Samples segregated by treatment (PBS vs PFF) on PC1, with additional genotype-dependent separation among PFF-treated mice on PC2. **(C)** Differential expression analysis and GSEA were performed for the indicated comparisons for the MSigDB Hallmark pathways, with genes ranked by log_2_FC × −log_10_(p-value). PFF-treated wild-type brains exhibited robust enrichment of inflammatory and immune pathways, including interferon-α and interferon-γ responses, allograft rejection, IL-2/STAT5 signaling, TNFα signaling via NF-κB, IL-6/JAK/STAT3 signaling, and complement activation. These pathways were markedly attenuated or failed to reach significance in Usp15^mut^ brains. Heatmap shows pathways reaching significance (adjusted p-value < 0.05) in any comparison, coloured by normalized enrichment score (NES). Grey indicates pathways that did not reach significance for the indicated comparison. **(D)** GSEA running enrichment graphs for the interferon alpha and interferon gamma response pathways comparing gene expression in *Usp15^mut^* vs WT brains treated with PFF.**(E)** Heatmap of leading-edge genes driving interferon-α and interferon-γ response signatures. Expression values are shown as log₂ fold change relative to genotype-matched PBS controls. PFF-injected wild-type mice exhibited strong induction of canonical interferon-stimulated genes (ISGs), including *Ifit1*, *Ifit3*, *Irf7*, *Mx1*, *Oas1*, *Oas2*, *Stat1*, *Psmb9*, and *Gbp2*, whereas *Usp15^mut^* brains showed markedly reduced ISG induction, with many transcripts remaining near baseline levels.

Further examination of the *Usp15*-dependent DEGs in PFF treated mice [(*Usp15^mut^* /PFF vs. *Usp15^mut^* /PBS) vs. (*Usp15*^+/+^ /PFF vs. *Usp15*^+/+^ /PBS) pair-wise comparisons] identified a significant effect of *Usp15* on expression of pro-inflammatory pathways. First, gene set enrichment analysis (GSEA) of MSigDB Hallmark pathways identified the same set of pathways activated in the brains of M83 mice, in both the WT and *Usp15^mut^* genotypes, following PFF injection (Figure 4C), similar to the PbA response (Figure 2C). Direct comparison of the *Usp15*^+/+^ and *Usp15^mut^* brains in the PFF condition, revealed a dampened activation of interferon-alpha and interferon-gamma responses, allograft rejection, fatty acid metabolism, and oxidative phosphorylation in the *Usp15* mutants (Figure 4C-D). Heatmap analysis of the DEGs in PFF-treated *Usp15^+/+^* vs *Usp15^mut^* mice highlighted the effect of *Usp15* inactivation on expression of ISGs, with lower activation in the *Usp15* mutants of several genes, particularly striking for *Ifit1*, *Ifit3*, *Irf7*, *Mx1*, *Oas1*, *Oas2*, *Stat1*, *Psmb9*, *Gbp2*, *Gbp3*, *Isg15*, *Usp18*, *Ifi44*, *Zbp1* and others (Figure 4E).

Hence ablation of *Usp15* dampens expression of neuroinflammatory pathways in the brain, in particular interferon-associated genes, following treatment with PFF but prior to appearance of clinical symptoms. This attenuated NI is associated with decreased progression of pathogenesis and neurodegeneration.

### *Usp15* knockdown suppresses type I Interferon pro-Inflammatory Programs in Activated Mouse Microglia

To further characterize Usp15-dependent cell identity programs, we performed GSEA using publicly available single-cell RNA-seq murine brain atlases as reference signatures, as previously applied in other neurodegeneration models^39^. This analysis revealed that, following PFF treatment, Usp15-deficient mice exhibited downregulation of microglia/macrophage and astrocyte signatures relative to WT, suggesting that Usp15 selectively modulates immune and astrocyte-mediated neuroinflammatory responses in the brain (Figure 5A).To explore the brain cell type(s) responsible for the *Usp15*-dependent effects on neuroinflammatory pathways detected in our in vivo model, we investigated the possible contribution of microglia and astrocytes in isolated primary cells. We purified primary neonatal mouse microglia and activated these cells with LPS. In parallel, we used siRNA-mediated treatment to silence USP15 expression prior to LPS treatment (Figure 5B). Both cell populations were then subjected to RNA sequencing, and DEGs were identified. This analysis revealed a strong enrichment of inflammatory gene signatures and ISG in LPS activated microglia with reduced expression in the siRNA treated group (Figure 5C). These results indicate that *Usp15* is required for full expression of inflammatory response pathways in primary microglia following activation by LPS. Using limma, we further compared the top DEGs from data sets obtained by comparing WT vs. *Usp15*/siRNA treated microglia with the top DEGs detected in the brains of WT and *Usp15* mutant M83 mice treated with PFF (see above and Figure 5C). The top ISGs showed a consistent pattern of downregulation upon Usp15 loss of function, with genes such as *Ifit3*, *Ifih1*, *Isg15*, *Gbp2, Ddx60*, *Irf7*, exhibiting strongly negative t-statistics in both the siRNA microglia dataset and the in vivo PFF conditions (Figure 5C). To directly compare the two perturbation approaches, we correlated limma t-statistics between the siRNA and knockout conditions and observed a positive correlation at both 48 and 72 hours post-siRNA (Figure 5D), indicating that knockdown of *Usp15* in microglia broadly phenocopies the genetic inactivation of USP15 in vivo in the PFF/M83 mouse model.

**Figure 5.**
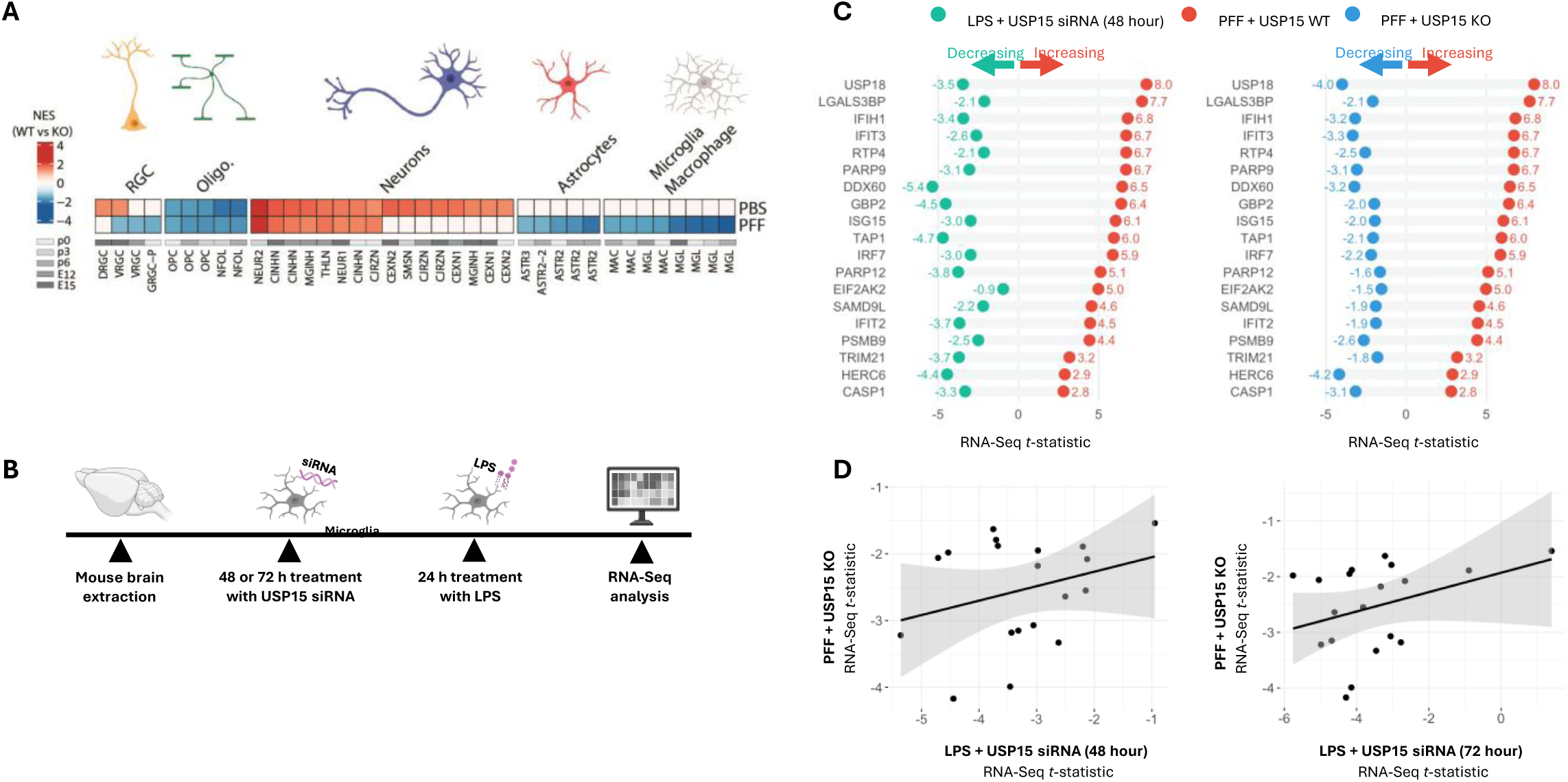
USP15 knockdown in microglia attenuates type I interferon and inflammatory gene expression. **(A)** Heatmap depicting normalized enrichment scores (NES) for WT vs. KO comparisons across major CNS cell-type lineages, in animals treated with PBS or PFF (rows). Columns represent individual gene signatures for radial glia cells (RGC), oligodendrocytes (Oligo.), neurons, astrocytes, and microglia/macrophages, corresponding to distinct embryonic and postnatal developmental stages (E12.5–P6, indicated by the greyscale bar). Color scale reflects NES, with red indicating positive enrichment and blue indicating negative enrichment in KO relative to the WT. test statistic. P-values were adjusted using the Benjamini-Hochberg procedure; signatures with adjusted p value < 0.05 were considered significantly enriched or depleted. **(B)** Schematic of experimental workflow. Primary microglia were isolated from neonatal mouse cortices, treated with USP15 siRNA for 48 or 72 h, stimulated with lipopolysaccharide (LPS) for 24 h, and subjected to RNA-seq analysis. **(C)** Differential gene expression analysis of interferon-stimulated and inflammatory genes in LPS-treated microglia transfected with Usp15 siRNA for 48 h (green, decreasing) compared to LPS alone or in bulk brain tissue from *Usp15* wild-type (WT) and *Usp15* knockout (KO) mice after α-synuclein preformed fibril (PFF) treatment (red, increasing and blue, decreasing, respectively). Data are presented as RNA-seq t-statistics for selected significantly altered genes. **(D)** Correlation of RNA-seq t-statistics between *Usp15* KO + PFF and *Usp15* siRNA + LPS conditions at 48 h (left) or 72 h (right) of siRNA treatment. Each point represents an individual gene. Solid lines denote linear regression fits; shaded areas indicate 95% confidence intervals.

Hence, Usp15 is a critical regulator of type I Interferon response in activated microglia. The concordance of *Usp15*-dependent DEG detected in vitro in activated microglia cells, and in vivo in the M83/PFF model further suggests that microglia are a strong contributor to *Usp15*-dependent effects in NI and neurodegeneration observed in vivo.

### USP15 Loss Functionally Blunts Type I Interferon Responses in Human iPSC-Derived Microglia

To investigate the role of USP15 in human primary microglia and primary astrocytes, we generated pluripotent human stem cells (iPSCs) from a control line (AIW002-2), in which USP15 had been inactivated by CRISPR-Cas9 targeting ^40, 41^. For this, we created two independent loss-of-function alleles, a complete USP15 knock-out null allele (USP15-KO) and a knock-in variant (USP15 L720R, referred to as USP15^mut^) corresponding to the human equivalent of the original mouse mutant (L749R) detected in our mouse genome screen for mutations protecting against NI ^33^.

Microglia were generated from these USP15 mutant lines using a stepwise hematopoietic specification protocol (Figure 6A; see Materials and Methods), including induction of hematopoietic progenitor cells, expansion of hematopoietic stem cells, and differentiation and maturation^40, 41, 42, 43^. This protocol afforded high reproducibility, enabling direct comparison of lineage integrity and inflammatory responses. USP15^KO^ and USP15^mut^ microglia displayed normal morphology and expressed canonical microglial markers (CD45, CD11b). Quantification of key microglial identity markers - Iba1, a cytoskeletal protein associated with microglial morphology and activation; PU.1, a transcription factor essential for microglial lineage specification; and TREM2, a receptor involved in lipid sensing and homeostatic function - confirmed expression of these markers across all three cell lines (Figure 6B). We further validated microglial commitment across genotypes, demonstrating intact expression of additional lineage markers (e.g., CX3CR1, P2RY12), robust phagocytic capacity, and preserved ramified morphology under basal conditions (Supplemental Figure 3). Together, these results indicate that USP15 loss does not alter microglial lineage commitment. Differentiated WT, USP15^KO^, and USP15^mut^ microglia were then activated with the viral double-stranded RNA mimic poly(I:C), and the effect of USP15 inactivation on the induction of ISG RNA was monitored by qPCR. WT microglia mounted a strong type I interferon response, with high induction of hallmark ISGs including MX1, MX2, IRF7, IFIT3, and USP18 (Figure 6C). In contrast, USP15^KO^ and USP15^mut^ microglia exhibited a markedly attenuated transcriptional response to poly(I:C), showing a 50–80% reduction in ISG induction relative to WT. These findings indicate that, alike mouse cells, USP15 is required for full activation of Type I Interferon transcriptional pathways in human microglia (Figure 5).

**Figure 6.**
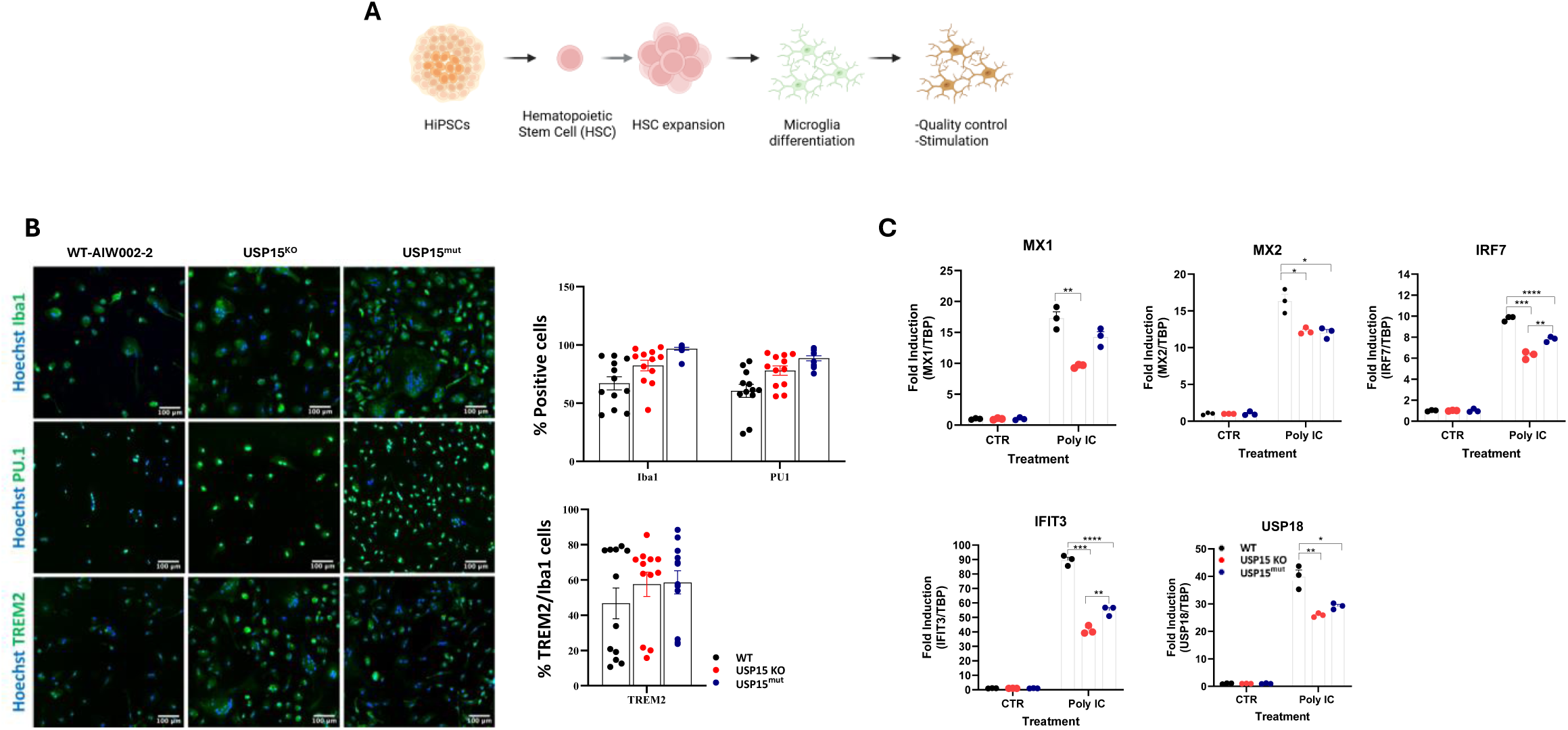
USP15 regulates interferon responsiveness in human iPSC-derived microglia. **(A)** Schematic representation of the work flow for the generation of microglia from human induced pluripotent stem cells (hiPSCs). **(B)** Immunofluorescence characterization of hiPSC-derived microglia. Representative images show expression of canonical microglial markers IBA1, PU.1, and TREM2 in wild-type (WT), USP15 knockout (USP15^KO^), and USP15^mut^ lines. Quantification shows the percentage of Iba1- and PU.1-positive cells (upper graph) and the percentage of TREM2-positive cells normalized to Iba1-positive cells (lower graph), indicating comparable differentiation efficiency across all three genotypes. Scale bars, 100 µm. **(C)** Interferon-stimulated gene (ISG) expression following poly(I:C) stimulation and measured by quantitative RT-PCR. Data are shown for the MX1, MX2, IRF7, IFIT3, and USP18 genes in either USP15^KO^ or USP15^mut^ microglia, and were conducted in triplicate. Statistical significance was assessed using appropriate multiple-comparison tests (*p* values as indicated).

### USP15 Loss of Function Suppresses Type I Interferon Responses in Human iPSC-Derived Astrocytes

We also generated human astrocytes from either WT or USP15-targeted iPSCs. Astrocyte production followed a protocol of derivation of neural progenitor cells (NPCs) and neurospheres, and serial cultures in neural induction media (Figure 7A; see Materials and Methods). NPCs were further differentiated into astrocytes by serial cultures in astrocyte differentiation medium, with astrocyte-specific markers expression confirmed by qPCR and immunofluorescence (S100β, GFAP)^44^.

**Figure 7.**
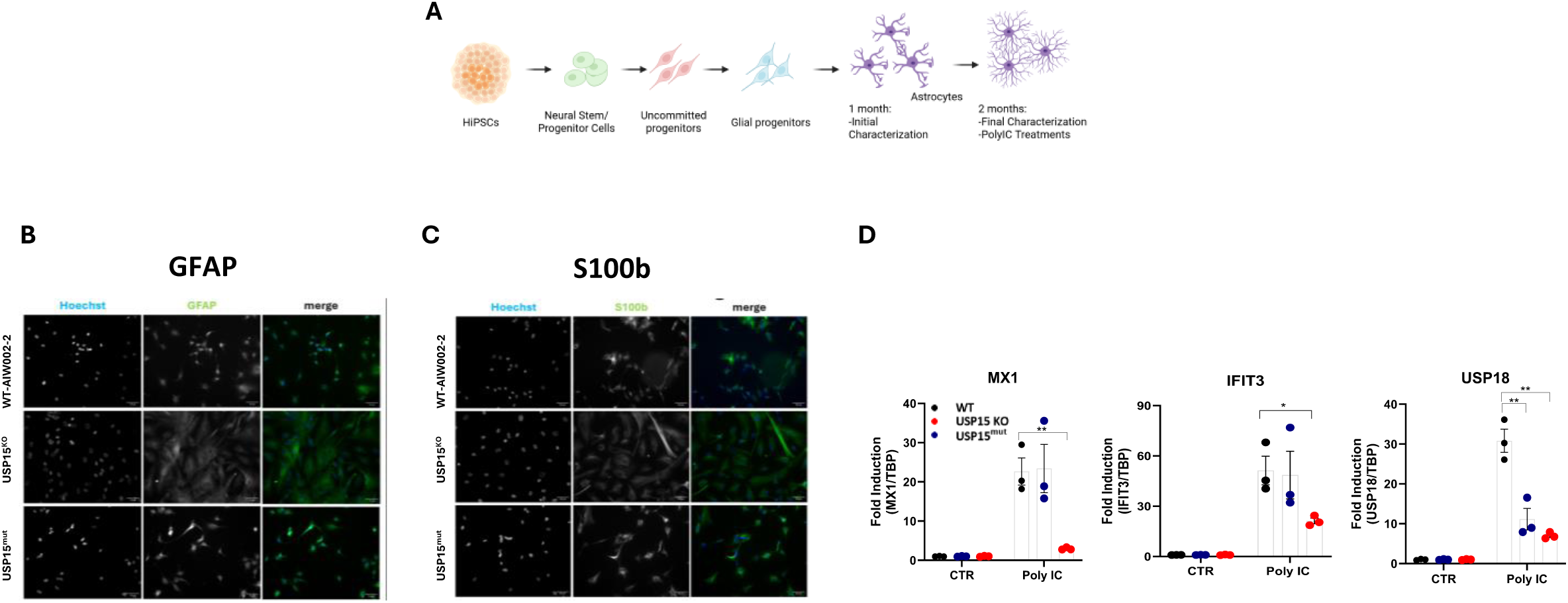
USP15 regulates interferon responsiveness in human iPSC-derived astrocytes. **(A)** Schematic representation of the work flow for generation of astrocytes from human induced pluripotent stem cells (hiPSCs). Initial characterization of the cells was performed at 1 month, while ISG response to poly(I:C) treatment was carried out at 2 months. **(B– C)** Immunofluorescence characterization of hiPSC-derived astrocytes. Representative images show expression of astrocytic markers GFAP (**B**) and S100β (**C**) in wild-type (WT), USP15 knockout (USP15^KO^), and USP15^mut^ astrocytes. Comparable morphology and expression of lineage markers was seen across genotypes. Scale bars as indicated. **(D)** Interferon-stimulated gene (ISG) expression following poly(I:C) stimulation and measured by quantitative RT-PCR. Data are shown for the MX1, IFIT3, and USP18 genes in either USP15^KO^ and USP15^mut^ astrocytes, and conducted in triplicate. Statistical significance was assessed using appropriate multiple-comparison tests (*p* values as indicated).

Immunostaining for canonical astrocyte markers revealed robust expression of GFAP (Figure 7B) and S100β (Figure 7C) across all genotypes. WT, USP15^KO^, and USP15^mut^ astrocytes displayed indistinguishable morphology, including elongated processes and typical astrocytic networks. USP15 loss does not impair astrocyte specification, maturation, or morphological differentiation, confirming that USP15 is not required for establishing human astrocyte identity. Mature astrocytes were then activated with the double-stranded RNA mimic poly(I:C), and the effect of USP15 deficiency on expression of pro-inflammatory response pathways was determined by qPCR analysis of indicator marker genes. WT astrocytes mounted a strong Type I interferon response, exhibiting robust induction of ISGs including MX2, and USP18 (Figure 7D). In contrast, USP15-deficient astrocytes showed a marked reduction in ISG induction.

These results in primary cells show that USP15 is required for full expression of pro-inflammatory pathways in general, and of type I IFN response ISGs in particular, in both human astrocytes and microglia in response to activation signals. These results further support the conclusion that both cell types contribute to the USP15-dependent transcriptional changes detected in the brain during NI, thereby directly contributing to the pathogenesis of neurodegeneration.

### USP15 is co-expressed with LRRK2 and SNCA in primary microglia from Parkinson’s disease patients

To further explore the neuroinflammatory relevance of USP15 in human PD, we examined its expression and co-expression patterns in publicly available single-cell RNA-seq data sets from PD patients’ microglia (Figure 8). USP15 was markedly enriched in PD microglia, while USP30, used here as a structurally and functionally related deubiquitinase control, showed no detectable expression in the same population (Figure 8). The majority of USP15-expressing microglia cells were also found to co-express LRRK2 (53.3%) and SNCA/α-synuclein (54.0%) (Supplementary Table 1), two proteins centrally implicated in PD pathogenesis^45^). These results indicate that in the microglial compartment USP15 is expressed and likely operates at the intersection of deubiquitination and PD-associated proteinopathy. Of additional relevance are recent large-scale plasma proteomic profiling studies that have identified USP15 as a consistently and strongly upregulated and differentially expressed protein in plasma from PD patients, coinciding with enrichment of neuroinflammatory pathways including TNF, NF-κB, and interferon α/β signaling^46^. These results, together with findings in wild-type and USP15 mutant human iPSC derived microglia and astrocytes (Figures 6,7), further support USP15 as a convergent node for NI and proteinopathy in human PD.

**Figure 8:**
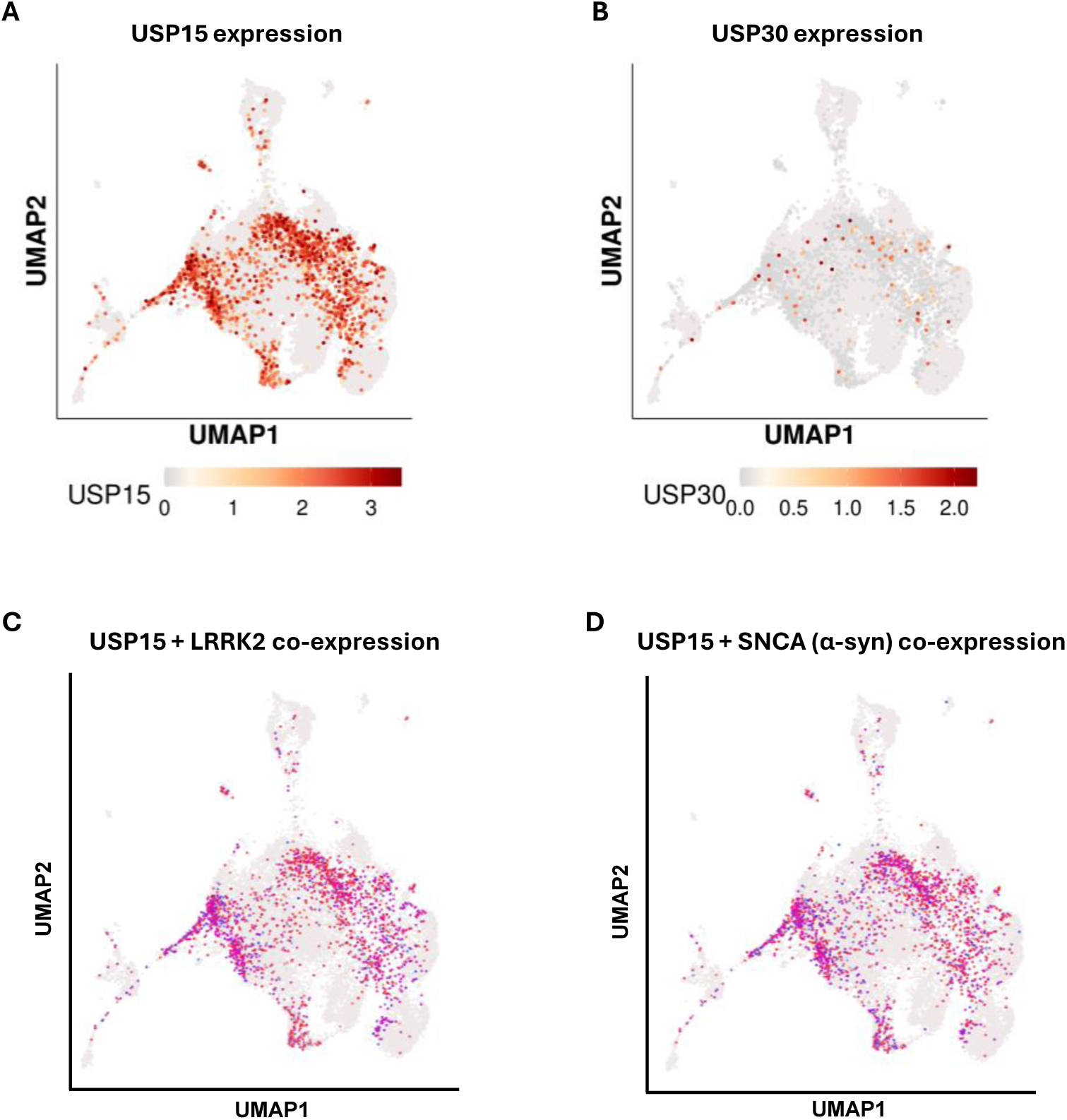
USP15 is co-expressed with LRRK2 and SNCA in primary microglia from Parkinson’s disease patients. Single-cell RNA-seq UMAP visualization and co-expression analysis of USP15 and USP30 in microglia from Parkinson’s disease (PD) patients. (**A**) USP15 is highly expressed and enriched in PD patient microglia. (**B**) USP30 shows no detectable expression in the same microglial population. (**C**) Co-expression analysis of USP15 and LRRK2: 53.3% of microglia co-express both genes, while 30.8% express USP15 alone, 7.5% express LRRK2 alone, and 8.4% express neither. (**D**) Co-expression analysis of USP15 and SNCA (α-synuclein): 54.0% of microglia co-express both genes, while 30.1% express USP15 alone, 7.7% express SNCA alone, and 8.2% express neither. Data derived from single-cell RNA-seq of PD patient microglia; source data accessed via ShinyCell Microglia (https://aysegulguvenek.shinyapps.io/shinyapp/).

### Genetic analyses of the USP15 locus on human Chromosome 12, and identification of a cis-acting regulatory eQTL

Human genetic data at the chr12q14.1 locus that contains *USP15* (with *TAFA2*, *MON2* and *PPM1H*) (Supplementary Figure 4) provide additional support for *USP15* as a candidate PD gene. The locus shows a suggestive PD-risk association in the largest European PD meta-GWAS to date with *P* = 3.0×10⁻⁶ and β = +0.095 for the lead variant rs4763185, and *USP15* receives a positive Polygenic Priority Score (+0.256) in the genome-wide gene-prioritization in that study. We undertook a GCTA-COJO analysis (a method that disentangles multiple independent genetic signals within a single genomic locus using GWAS summary statistics) and showed that the PD signal at the locus decomposes into two statistically independent components (marked as gold stars in Supplementary Figure 4) at rs4763185 (4.6 kb 5’ of *PPM1H*) and rs11174279 (intronic in *TAFA2*).

We examined the effect of these and other neighbouring genetic variants on the expression of the four genes in monocytes using publicly available eQTL data from Quach et al. (n = 200 evaluated under five conditions^47^). *USP15* and *TAFA2*, but not *MON2* nor *PPM1H*, show monocyte cis-eQTL signals concentrated in the chr12:62.1–62.4 Mb interval where the two genes overlap (Figure 9). Bayesian colocalization showed shared causal variation between the PD GWAS signal and the monocyte cis-eQTLs of both USP15 and TAFA2 (PP.H4 ≈ 0.81 for both genes). Conditional analysis identified a single haplotype (∼0.03 frequency in Europeans) carrying the principal cis-eQTL associated variants within and the PD-tagging risk variant at rs4763185 in coupling-phase LD (|D′| = 0.82, r² = 0.08), while the second signal at rs11174279 is on a separate axis of PD risk that is independent of the eQTL (Supplementary Figure 5). Independent confirmatory evidence was obtained in CD14+ monocytes from the BLUEPRINT study (Figure 10; n = 191^48^), and in a smaller series from the DICE study (see Supplementary Note 1; n=91^49^).

**Figure 9:**
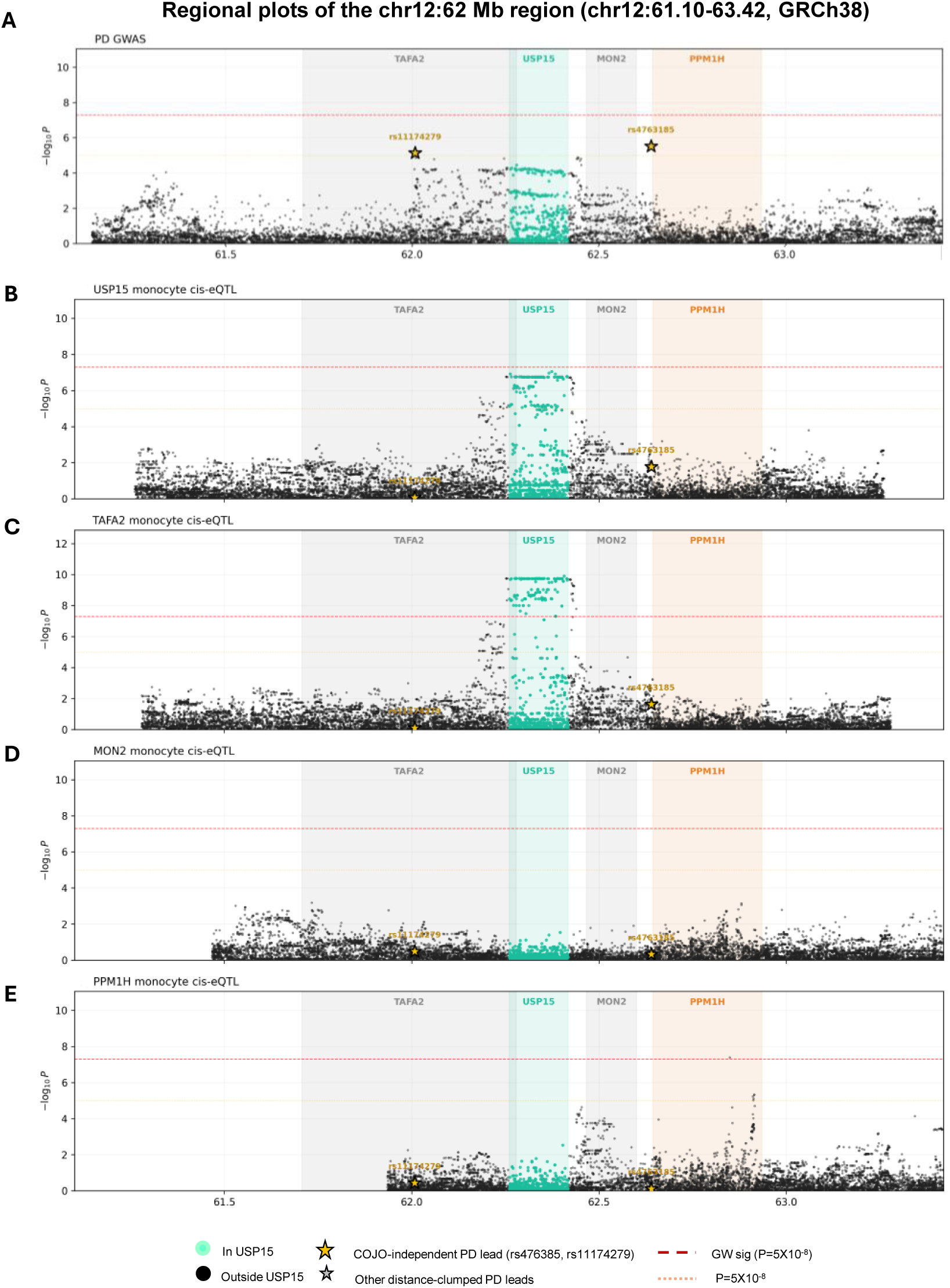
Genetic analyses of the USP15 locus on human Chromosome 12 and identification of a cis-acting eQTL. **(A)** Manhattan plot for genome wide association (-log_10_P) of SNPs from the *USP15* locus on 12q14.1 (European PD meta-GWAS analysis), including positions of the *TAFA2* (blue), *USP15* (green), *MON2* (grey), and *PPMH1* (orange) genes and top associated SNPs, rs11174279 and rs4763185 (P=5 x 10^−6^; shown as gold stars). Manhattan plots showing the association of SNPs from the region with expression levels of **(B)** *TAFA2*, **(C)** *USP15*, **(D)** *MON2* and **(E)** *PPMH1* mRNAs in a cohort of CD14+ monocytes from primary cis-eQTL panel of Quach et al, ^47^, and identifying a highly significant cis-eQTL overlapping the *TAFA2* and *USP15* genes. The strength of the association (-log_10_P) and significance threshold (orange stippled line) are indicated.

**Figure 10:**
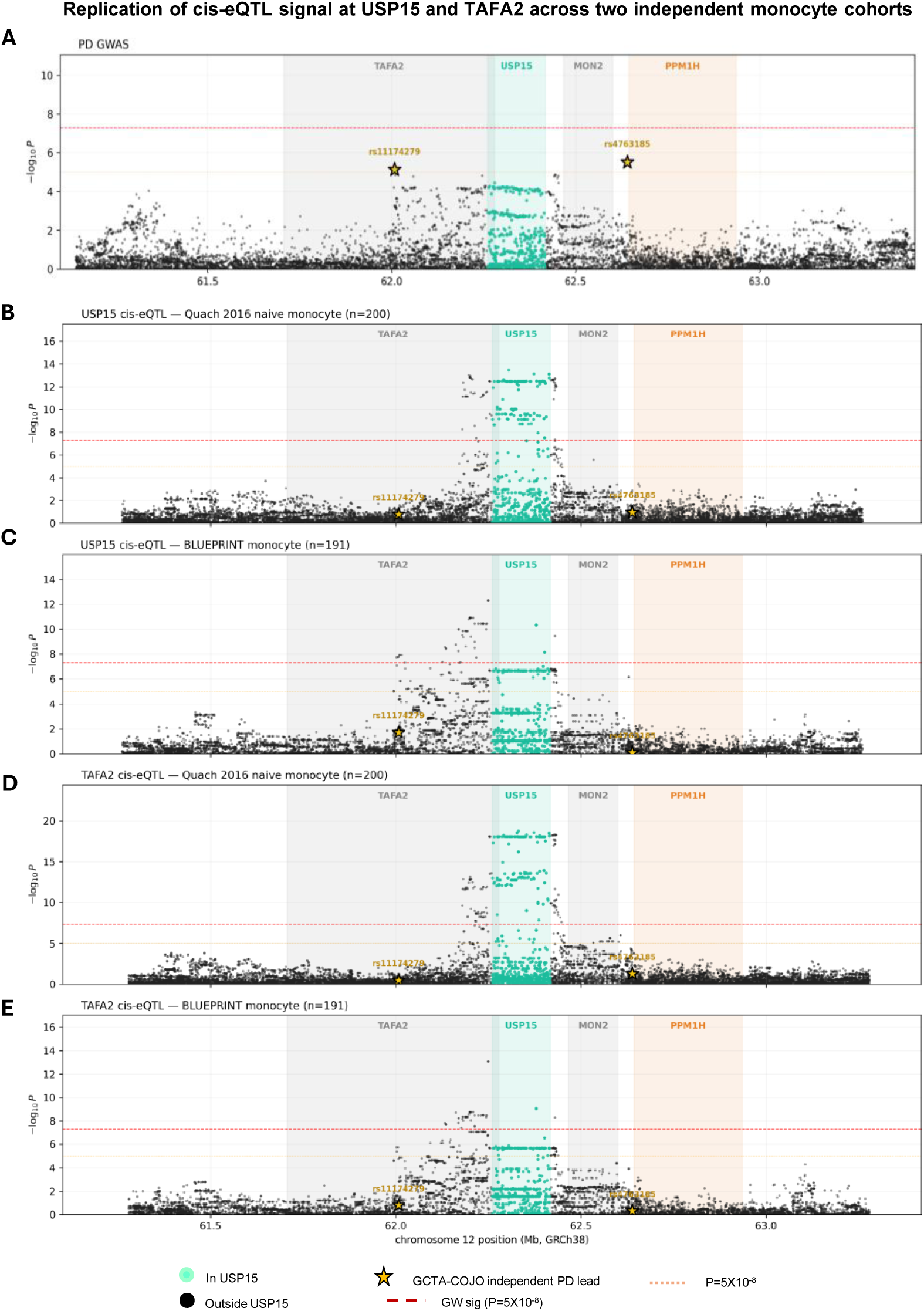
Validation of a cis-acting eQTL at the USP15 locus in different monocyte cohorts. **(A)** Manhattan plot for genome wide association (-log_10_P) of SNPs from the USP15 locus on 12q14.1 (European PD meta-GWAS analysis) as in legend to Figure 9. Manhattan plots showing the association of SNPs from the region with expression levels of (**B,C**) *USP15* and (**D,E**) *TAFA2*in independent cohorts of naïve CD14+ monocytes from Quach et al, ^47^ (**B,D**) and from BLUEPRINT ^53^ (**C,E**). The strength of the association (-log_10_P) and significance threshold (orange stippled line) are indicated.

Two-sample Mendelian randomization with cis-eQTL instruments showed that increased monocyte expression is significantly associated with increased PD risk (pleiotropy-robust MR-link-2 α = +0.022, P = 6.1×10⁻⁶). Decomposing the joint signal further, the haplotype carrying the cis-eQTL elevating G allele at rs7309209 but not the rs4763185-A risk allele (the GG haplotype, ∼18% frequency in Europeans) returns a per-chromosome OR of 1.025 (95% CI 1.007-1.043, P = 5.6×10⁻³) showing that the cis-eQTL allele itself confers a small but nominally significant PD-risk increase independent of rs4763185 (Supplementary Figure 5). Further details of the statistical analysis are provided in Supplementary Note 1.

Together, these analyses strongly suggest that USP15 and its associated cis-eQTLs in myeloid cells are linked to PD pathogenesis; These findings support our data in mouse experimental models of PD, and establish USP15 as a therapeutic target for drug discovery in PD.

## DISCUSSION

Using an unbiased genome-wide in vivo screen for genes and pathways driving pathological NI, we have identified USP15 as an important regulator. USP15 inactivation protects mice against lethal NI in acute microbial (experimental cerebral malaria; ECM) and autoimmune (experimental autoimmune encephalomyelitis; EAE) models ^33^. Time course studies with bulk RNA sequencing during pre-symptomatic phase of ECM showed overall dampened expression of inflammatory pathways (GSEA signatures) seemingly contributed by both resident brain cells early during pathogenesis (innate immune responses, Type I and type II IFN and TLR associated signatures, ISGs) with additional contributions concomitant to the infiltration activated myeloid cells and activated lymphoid cells (myeloid and lymphoid specific signatures) in the brain. These results, together with the observations that a) USP15 is expressed in all peripheral lymphoid and myeloid subsets, and in both astrocytes and microglia, and b) that differential ECM resistance/susceptibility associated with the mutant or WT USP15 genotypes, respectively, can be transferred in reciprocal bone marrow chimeras, indicated a functional role of USP15 in both resident brain cells and peripheral leukocytes.

We herein conducted several in vivo and in vitro experiments to further probe the role of USP15 in modulating pro-inflammatory functions of resident brain astrocytes and microglia *in situ*. Studies by bulk RNA sequencing in brains of mice undergoing ECM and specifically ablated for USP15 in astrocytes and microglia identified Type I and Type II interferon pathways (and Interferon-stimulated genes; ISGs) as the dominant and most significantly affected by the loss of USP15 (Figure 2C,2D). This signature of differential expression of ISGs was validated in studies of primary iPSC-derived human astrocytes and microglia targeted for USP15 inactivation following activation with viral mimics (Poly:IC)(Figure 7D). The same Type I and Type II interferon and ISGs pathways were differentially expressed in both primary mouse microglia defective in USP15 (siRNA) and activated with LPS (Figure 5B). Together, these results indicate that resident microglia and astrocytes phenotypically express the USP15 genetic effect on NI in the ECM model through differential expression of Type I/II IFNs and associated ISGs.

In parallel, we observed that inactivation of USP15 protects mice against a-synucleinopathies in the M83/PFF model, with dampened pathogenesis, delayed disease progression and increased survival (Figure 3). RNA-seq analyses of contra-lateral brain prior to the appearance of neurological symptoms again identified differential expression of the same USP15-dependent Type I/II IFNs and ISGs pathways (Figure 4) previously detected in a) total brain and spinal cord of WT and USP15 mice undergoing ECM and EAE, respectively^33^; b) total brain from WT and mutant mice conditionally deleted for USP15 in microglia and undergoing ECM (Figure 2); c) primary mouse microglia ablated or not for USP15 (siRNA), and activated with LPS (Figure 5). In addition, this USP15 signature was also detected in brain tissues of TRIM25 mutant mice undergoing ECM ^33^. We have previously demonstrated that USP15 genetically and physically interact with TRIM25, causing the de-ubiquitination of TRIM25 and consequent activation of its E3 ligase activity ^33^ to ubiquitinate and stabilize the CARD domains of RIG-I and activate the RIG-I signaling complex to induce Type I IFN in response to viral ds-RNA and other PAMPs ^34, 35^. Hence, the identification of Type I IFN and ISGs pathways as the shared and dominant component of USP15 and TRIM25 signatures is completely compatible with the known role and molecular mechanism of action of these 2 proteins in Type I IFN signaling. In agreement with these results, we recently observed that TRIM25 defective mice are also protected against synucleinopathies in the M83/PFF model, with reduced pathogenesis and increased survival time (data not shown).

How relevant are these findings in mice to human Parkinson’s Disease? Several lines of evidence reported here indeed suggest that USP15 may play a role in PD pathogenesis. Firstly, we show that inactivation of USP15 in primary iPSC-derived human microglia and astrocytes impairs pro-inflammatory functions of these cells, notably activation of Type I IFN and ISG responses (Figures 6 and 7). These responses are known to promote NI, contributing to pathogenesis in PD patients brains *in situ* ^50^. Secondly, our analysis of public single cell RNA-seq datasets shows that USP15 is strongly expressed in a subset of human microglia from PD patients, where it is co-expressed with LRRK2 and SNCA (Figure 8), two proteins that play a direct role in onset and progression of PD ^51, 52^. Thirdly, a recent study, by Cha et al. implicates USP15 as a relevant protein in human PD ^46^. In large-scale proteomic studies of plasma proteins (DEPs) from PD patients and controls (Global Neurological Proteomics Consortium; 462 PD cases, 1186 controls), they used unsupervised clustering analysis to identify disease sub-types, map differentially expressed plasma proteins (DEPs) and associated biological pathways. This analysis identified 2 PD sub-types, Type I and Type II, which differ in extent of NI, oxidative stress and mitochondrial programs; Type II PD was characterized by enrichment of inflammatory and stress-response proteins. Notably, USP15 was strongly and significantly upregulated in both PD sub-types but was particularly elevated in Type II PD patients plasma. In addition, GSEA analysis identified “de-ubiquitination” as the most enriched pathway for both Type I and Type II PD, consistent with increased plasma overexpression of USP15. The co-enrichment of USP15 and its de-ubiquitinase activity in both Type I and Type II PD plasma samples strongly supports a role in pathogenic mechanisms of human PD. Fourthly, genetic and epigenetic studies of human USP15 further support a possible role in PD. The chr12q14.1 locus that contains USP15 shows suggestive evidence of association with PD risk in the largest European meta-GWAS to date (lead rs4763185, *P* = 3.0×10⁻⁶;^36^), with *USP15* receiving a positive Polygenic Priority Score in the same study’s gene-prioritization analysis. However, this evidence is reinforced by the additional analyses presented here: cis-eQTL colocalization in primary monocytes (PP.H4 ≈ 0.81 for *USP15 and TAFA2*), a positive Mendelian-randomization estimate of cis-regulated *USP15* expression on PD risk that survives pleiotropy-robust cis-MR (MR-link-2 α = +0.022, *P* = 6.1×10⁻⁶) ^36^, and concordant regulatory effects across independent monocyte cohorts (BLUEPRINT replication PP.H4 = 0.80). Because *USP15* and *TAFA2* share most of their cis-eQTL architecture, the monocyte regulatory signal implicates this two-gene USP15/TAFA2 cis-window collectively. Crucially, within this shared window, the plasma proteomic data of Cha et al. distinguish the two genes: *USP15*, but not *TAFA2*, is overexpressed in PD patient plasma ^46^. The studies of USP15 in mouse models in vivo complemented by cellular and molecular data presented elsewhere in this work, together strongly suggest that USP15 is the specific causal gene at this locus.

Together, our findings identify USP15 as a strong, biologically supported candidate for drug discovery and therapeutic intervention in PD.

## Acknowledgements

This work was supported by research grants to PG from the Consortium Quebecois de la Recherche sur le Médicament, Corbin Therapeutics, Canada First Research Excellence Fund (HBHL and D2R programs), and from the Michael J Fox Foundation. This work was supported in part by the Alzheimer’s Disease Data Initiative, California Office of the Small Business Advocate and Immuneering (Alleo). Julia Luo is supported by a Vanier Canada Graduate Scholarship. Étienne Collette was supported by a Fonds de la Recherche Québec-Nature et Technologie (FRQ-NT) master’s scholarship. David Langlais is supported by a Fonds de la Recherche Québec-Santé (FRQ-S) Chercheur-Boursier Junior 2 Award (#348782) and the Canada Foundation for Innovation John R. Evans Leaders Fund (#38033). We thank Andrea Krahn, Julien Sirois, Wolfgang Reintsch, Wen Luo, and Genevieve Dorval from the Neuro’s Early Drug Discovery Unit (EDDU) for technical support during the project.

## Author contributions

NF, YC, ML, EDC-P, VP,JR, MK: Study design, experimental work, data analysis and manuscript writing. PG, DL, EF, TD: Study design, data analysis and manuscript writing. JXXL, MW, J-FO: Experimental work and data analysis. AA, IS, ZY, CX-QC, NA, JJM: Experimental work. AA, EC, NJ, CLK data analysis.

## Competing Interests

The authors declare no competing interests.

---

**Supplementary Figure 1. Differential gene expression analysis of *Plasmodium berghei*–infected brains reveals cell-specific effects of *Usp15* deficiency in astrocytes and microglia.**

Heatmap of interferon alpha and interferon gamma response genes significantly regulated in any comparison. Expression values are shown as log₂ fold change relative to genotype-matched PBS controls. PbA-infected WT brains exhibited strong induction of classical interferon-stimulated genes (ISGs), including *Ifit3*, *Irf7*, *Mx1*, *Oas2*, and *Stat1*.

**Supplementary Figure 2. USP15 selectively amplifies interferon-driven neuroinflammatory gene programs induced by α-syn pathology.**

**(A)** Volcano plots highlighting DEGs due to infection (first row) and genotype effects in the context of infection (second row). Red indicates significantly differentially expressed genes at adjusted p-value < 0.05, absolute log2FC > 1. **(B)** Venn diagram illustrating the overlap of PFF-induced DEGs between WT (WT PFF vs WT PBS) and *Usp15^mut^* (*Usp15^mut^* PFF vs *Usp15^mut^* PBS) brains. A substantial subset of genes was shared between genotypes, while distinct DEG sets were uniquely regulated in WT or *Usp15^mut^* mice. **(C)** Heatmap of all significantly PFF-induced DEGs, grouped according to whether genes were regulated in both genotypes, exclusively in WT, or exclusively in *Usp15^mut^* mice. Samples are shown by treatment (PBS vs PFF) and genotype (WT vs *Usp15^mut^*). Heatmap values represent log₂ fold change relative to genotype-matched PBS averages. For each DEG group, over-representation analysis was performed using MSigDB Hallmark pathways, with significantly enriched pathways (adjusted p value < 0.05) indicated. Genes induced exclusively in WT brains were enriched for interferon and inflammatory signaling pathways, whereas shared DEGs reflected USP15-independent responses to α-syn pathology.

**Supplementary Figure 3: USP15 loss does not impair phagocytic activity in human iPSC-derived microglia.**

(**A**) Phagocytic activity of hiPSC-derived microglia (iMGL) across genotypes. Representative fluorescence images show uptake of Zymosan particles (red) counterstained with Hoechst (blue) in wild-type (WT), USP15^KO^, and USP15^mut^ microglia under four conditions: unstimulated (CTR), LPS (24h), Pam3CSK5, and IFNγ + TNFα. Quantification of the percentage of phagocytic cells is shown for each genotype and stimulation condition. No significant difference in phagocytic capacity was observed across genotypes under any condition tested. (**B**) Phagocytic activity restricted to TREM2-positive microglia. Representative images show uptake of Zymosan particles (red) in TREM2-positive cells (green), counterstained with Hoechst (blue), across the same genotypes and stimulation conditions as above. Quantification shows the percentage of TREM2-positive cells with phagocytic activity. No significant difference was detected across genotypes, confirming that USP15 loss does not impair the phagocytic function of homeostatic TREM2-expressing microglia. Scale bars as indicated. Data are presented as mean ± SEM.

**Supplementary Figure 4: Genomic organization of the USP15 locus on human Chromosome 12q14.1.**

The position of the TAFA2 (blue), USP15 (green), MON2 (grey), and PPMH1 (orange) genes on Chromosome 12q14.1 is indicated, together with the size of the genes (in kb) and the direction of transcription (white arrow), with the area of transcriptional overlap between TAF2 and USP15 shown (yellow box). The position of informative SNPs, rs11174279 and rs4763185 used for PD risk association, and haplotype mapping are indicated. The locus coordinates on GRCh38 are indicated in Mb (bottom scale).

**Supplementary Figure 5: Haplotype structure of the USP15 locus on human Chromosome 12q14.1.** (**A**) Linkage disequilibrium structure among the PD-associated risk and the cis-active eQTL active markers. (**B**) The two-locus haplotype frequencies based on genotypes at rs4763185 and rs7309209 are shown. Five-marker linkage-disequilibrium analysis among the two GCTA-COJO independent PD leads (rs11174279, rs4763185; outlined in red) and the three cis-eQTL-active variants in the *USP15* / *TAFA2* shared cis-window (rs7307385, rs12311279, rs7309209; outlined in teal). All values are computed from UK Biobank chr12 imputed phased haplotype dosages restricted to *n* = 339,334 unrelated European-ancestry individuals (*n* = 678,668 chromosomes). (a) Pairwise *r²* heatmap. (b) Pairwise |*D*′| heatmap; |*D*′| is the appropriate metric for the haplotypic interpretation when one variant has very skewed allele frequencies, because it normalizes out asymmetric MAF. The contrast between panels (a) and (b) at the rs4763185 row/column reflects the low frequency of the rs4763185 risk allele (0.030): *r²* is capped well below 1 even when |*D*′| is high. (c) Empirical chromosome-frequency table for the two-locus haplotype rs4763185 × rs7309209, counted directly from phased imputed haplotype dosages without any model-based phase imputation, over the 336,795 individuals (673,590 chromosomes) with non-missing imputed phased calls at both loci. The teal-highlighted bar shows the AG haplotype carrying the PD-risk A allele at rs4763185 together with the *USP15* cis-eQTL elevating G allele at rs7309209; it is rare (0.026 of EUR chromosomes; *n* = 17,327 of 673,590) but is approximately 4-fold enriched over the frequency expected under independence. Inset summary statistics: *r²* = 0.080, |*D*′| = 0.823 (coupling), *f*(rs4763185 A, PD risk) = 0.030, *f*(rs7309209 G, USP15 cis-eQTL elevating) = 0.207.

## MATERIALS AND METHODS

### Animals and Ethical Approval

All studies in mice were conducted in accordance with institutional and national guidelines of the Canadian Council on Animal Care and all research protocols and ethics were approved by the McGill University Animal Care Committee. Mice were maintained in a specific pathogen-free facility on a 12 h light/dark cycle with ad libitum access to food and water.

### Generation of Conditional USP15 Knockout Mice

USP15-tm1c [C57BL / 6N-Usp15tm1c(EUCOMM)Wtsi / Tcp] (USP15^flox/flox^) mice were generated at the Centre for Phenogenomics of the University of Toronto from a frozen stock initially obtained from the European Conditional Mouse Mutagenesis program. The original line was expanded by crossing to C57BL/6J mice and intercrossed to generate *Usp15^flox/flox^* homozygotes. These were then backcrossed to either Aldh1l1-CreERT2 mice to generate astrocyte-specific USP15 knockouts or to Cx3cr1-CreERT2 mice to generate microglia-specific *Usp15* knockouts. Genotypes were confirmed by PCR using DNA extracted from tail biopsies. Targeted excision of the USP15 gene was induced in adult mice using tamoxifen (100 μg/g, intraperitoneal, administered daily for 5 consecutive days).

### Plasmodium berghei ANKA Infection Model

Phenotyping for susceptibility to experimental cerebral malaria using infection with *Plasmodium berghei* ANKA was carried out as we previously described ^33^. Tamoxifen-treated *USP15^flox/flox^*-Cre transgenic mice and control littermate lacking the Cre transgene were infected intraperitoneally with freshly isolated 10^6^ *P. berghei* ANKA-parasitized erythrocytes two weeks after the last tamoxifen injection. This two-week interval allows peripheral monocytes to repopulate from *Cx3cr1*-negative bone marrow progenitors, while brain microglia retain the recombination ^54^. Animals were monitored daily for the appearance of clinical symptoms (ruffled fur, hunched back, lethargy, weight loss, partial paralysis) and for survival according to pre-determined humane end-points. For RNA sequencing, mice were anesthetized with 2% isoflurane, perfused with PBS, and brains were harvested on day 5 post-infection, prior to the appearance of clinical signs, and flash-frozen for future use.

### Generation of TgM83^+/−^/Usp15^mut^ mice

Usp15^L749R^ homozygous mutant mice (namely *Usp15^mut^*) were initially generated and identified in a whole genome N-ethyl-N-nitrosourea (ENU) mutagenesis scan, looking for genes whose inactivation protects against P. berghei ANKA-induced cerebral malaria ^33^. The ENU-induced mutant *USP15* allele (*Usp15^L749R^*, *Usp15^mut^*) was identified in a multi-generation breeding scheme and subsequently backcrossed to homozygosity onto the C57BL/6J background for four generations before establishing the homozygous Usp15^L749R^ line by intercrossing.

Hemizygous TgM83^+/−^ mice (B6;C3H-Tg[SNCA]83Vle/J; JAX strain #004479) were bred in-house. Hemizygous M83 mice (M83^+/−^), express multiple copies of human alpha-synuclein bearing the familial PD-related A53T mutation under the control of the mouse prion protein promoter ^37^. Hemizygous TgM83^+/−^ and *Usp15^mut^* mice were intercrossed to generate TgM83^+/−^ *Usp15^mut^* and wild-type littermates M83^+/−^ *Usp15^+/+^*. Animals were maintained at the Montreal Neurological Institute (The Neuro) animal facility at McGill University under standard laboratory conditions.

### Induction of α-synucleinopathy in mice

Three-month-old mice (both females and males) were housed in individual cages and pre-treated with carprofen (20 mg/kg) and bupivacaine (25 mg/kg) before surgery for multimodal analgesia. Mice were anesthetized with 2% isoflurane and stereotactically injected in the right dorsal striatum with either PBS or pre-formed α-synuclein fibrils (PFFs; 12.5 μg per brain) using a 5 μL Hamilton syringe fitted with a 33-gauge needle, at the following coordinates: +0.2 mm relative to bregma, +2.0 mm from midline and 2.6 mm depth below the dura, as previously described ^38, 55^. All mice were injected on the same day and randomly assigned to two experimental groups.

The first group (endpoint cohort) was used for RNA-seq and immunohistochemistry analyses. Mice were euthanized at the experimental endpoint, when the first mouse reached a humane endpoint (day 84 post-injection). All mice in the group were euthanized simultaneously. For tissue collection, mice were anesthetized with 2% isoflurane and perfused with PBS alone (RNA-seq) or PBS followed by 10% formalin (immunohistochemistry). The second group (survival cohort) was used to monitor disease progression and survival. Mice were weighed and assessed regularly for clinical signs. Humane endpoints were defined as severe motor impairment preventing access to food or water, or body weight loss exceeding 20%. In this cohort, each mouse was euthanized individually upon reaching its humane endpoint.

### Immunohistochemistry

The brains were removed and fixed in 10% formalin for 24 h at 4°C, then incubated in 70% ethanol for 7 days and processed for paraffin embedding. Coronal brain sections were cut with a paraffin microtome at 5 μm thickness. Tissue sections were incubated in citrate buffer (pH 6.0) for 10 min, rinsed with Tris-buffered saline containing 0.1% Tween-20 (TBST), and incubated in 3% hydrogen peroxide for 15 min. Subsequently, the sections were blocked with 10% normal goat serum in TBST for 30 min at room temperature and incubated with anti-phospho-syn antibody (1:500, Abcam, ab184674) overnight at 4°C. The next day, sections were incubated with horseradish peroxidase (HRP)-conjugated secondary antibodies (1:500, Jackson ImmunoResearch) for 30 min. The peroxidase reaction product was visualized as a brown precipitate by incubating the tissue with the DAB substrate kit (cat.#8059, Cell Signaling Technology). Sections were counterstained with hematoxylin to visualize nuclei. Three coronal sections containing the substantia nigra (minimum inter-section distance sections 40 μm), were examined by a bright-field microscope using an Olympus DP-21SAL coupled to a DP21/DP26 digital camera. The images were analyzed using Fiji-ImageJ 1.53 software. Macros for Fiji-ImageJ were written in Jython using basic ImageJ functions. Each spot labeled with pSyn was defined as a as a pSyn positive particle, characterized by perimeter, area, and size.

### Bulk RNA Sequencing on Brain tissues and analysis

Bulk RNA-seq of total brain tissue was performed as previously described ^33^. Brains were stored at −80°C and homogenized in TRIzol reagent to extract RNA. RNA integrity was validated (RIN > 8) prior to library preparation. Libraries were prepared using poly(A)-enriched RNA-seq kits. Libraries were sequenced on an Illumina NovaSeq 6000 platform in a paired-end 100bp configuration. Adapter sequences and low-quality bases (Phred score < 30) were trimmed from reads using Trimmomatic v0.39 ^56^. The resulting reads were aligned to the *Mus musculus* mm10 reference genome, using HISAT2 v2.2.1 ^57^. Expression levels were quantified by strand-aware counting of reads aligning to exonic features using featureCounts (Subread package version 2.0.3) ^58^ with annotations from GRCm38. Raw read counts were filtered to remove residual rRNA and mitochondrial reads, and only genes with an expression level above a threshold of 10 counts per million reads (CPM) in at least 3 libraries were retained. A principal component analysis (PCA) was computed on the CPM values. Filtered CPMs were normalized using TMM normalization and then fitted to a genewise negative binomial generalized linear model using the EdgeR package version v3.42.4 (Robinson et al, 2010). Differential gene expression analysis was performed to assess the impact of the infection across the genotypes. Significantly dysregulated genes were selected using a log2 fold change threshold of >|1| and an FDR < 0.05 (Benjamini–Hochberg corrected *p* values). Gene set enrichment analyses were performed using the clusterProfiler Bioconductor package ^59^ with the msigdbr database.

### Gene set enrichement analysis for cell type signatures

Raw reads were trimmed and illumina specific sequences were removed using Trimmomatic v0.32. A four-nucleotide sliding window was used to remove the bases once the average quality within the window fell below 30 and reads shorter than 30 base pairs were dropped. Reads were aligned to the mouse reference genome build mm10 using STAR v2.3.0e with default settings and and expression levels per gene were estimated by quantifying reads uniquely mapped to exonic regions defined by ensGene annotation set from Ensembl (GRCm38, n=39,017 genes) using featureCounts (v1.4.4). Normalization (mean-of-ratios), variance stabilized transformation of the data and differential gene expression analysis were performed using DESeq2 (v1.14.1) for WT vs Usp15 KO for PBS and PFF seperatly.

To evaluate cell type composition in the bulk RNA seq samples, a reference panel of 82 gene signatures covering the major cell type linages in brain, was used as input for gene set enrichment analysis (GSEA)^60^. GSEA was run using the GSEA function from clusterProfiler (v4.14.6), ranking the genes by the Negative Binomial Wald test statistic. P-values were adjusted using the Benjamini-Hochberg procedure; signatures with adjusted p value < 0.05 were considered significantly enriched or depleted.

### Preparation and Culture of Primary Mouse Microglia

Primary microglia were prepared from postnatal day 0–3 C57BL/6J mouse pups. Brains were removed and cortices dissected in calcium- and magnesium-free Hank’s Balanced Salt Solution (HBSS) on ice. Cortical tissue was minced with a sterile razor blade and digested in 0.25% trypsin in HBSS (20 min, 37°C), before terminating digestion with a stop solution containing 0.1% DNase I and 0.5% trypsin inhibitor in fetal calf serum (FCS). Tissue was triturated 10–15 times through a fire-polished Pasteur pipette and centrifuged at 300 g for 5 min. Pellets were resuspended in Dulbecco’s Modified Eagle Medium (DMEM) supplemented with 10% FCS, 1% penicillin-streptomycin, and 2 mM L-glutamine, at a final volume of 1 mL/pup. Cells were seeded into poly-D-lysine-coated T75 flasks containing 14 mL of nutrition medium (1 mL/pup) and maintained at 37°C, 5% CO₂, with medium changes every 3–4 days. After 14 d in vitro, microglia were harvested by shaking flasks at 200 rpm for 2 h at 37°C. Detached cells were collected, centrifuged at 300 g for 10 min, resuspended in nutrition medium, and plated in uncoated 48-well plates (1.5 × 10⁵ cells/well). After 2 h at 37°C, plates were rocked at 100 rpm for 5 min to dislodge non-microglia, and the medium was replaced to on day in vitro (DIV) 2 medium. Microglia were incubated for 24 h before siRNA treatment.

### siRNA Transfection and LPS Stimulation for Primary Mouse Microglia

On day in vitro (DIV) 2, medium was replaced with serum-free Accell Delivery Medium (Dharmacon, B-005000). *Usp15* Accell siRNA (mouse, catalog number: E-040417-00-0005; Dharmacon) was reconstituted to 100 μM in 1× siRNA buffer (Dharmacon) and incubated for 70–90 min at 37°C. A 1 μM delivery mix was prepared by adding 3 μL of siRNA stock to 297 μL Accell Delivery Medium. Culture medium was replaced with 300 μL/well of delivery mix, and cells were incubated for 48 or 72 h. For LPS stimulation, medium was replaced with serum-reduced DMEM (5% FCS, 2 mM L-glutamine) and supplemented with 100 ng/mL lipopolysaccharide (LPS; Sigma, L6529) for 24 h. Control wells received identical treatment without LPS. Cells were lysed in TriFast (peqGOLD, 30-2030) for RNA analysis.

### RNA Extraction and RNA-seq for Primary Mouse Microglia

RNA was extracted using the peqGOLD TriFast kit according to the manufacturer’s instructions. Cells were lysed in 500 μL TriFast per well, followed by addition of 100 μL chloroform, vortexing, and phase separation (12,000 × g, 5 min, RT). The aqueous phase was mixed with 100 μL of 2-propanol, incubated on ice for 5–15 min, and centrifuged (12,000 × g, 10 min, 4°C). Pellets were washed twice with 75% ethanol, air-dried, and resuspended in RNase-free water. RNA quantity and purity were determined using a NanoDrop 1000 spectrophotometer (Thermo Fisher Scientific). One microgram of total RNA was reverse transcribed using the QuantiTect Reverse Transcription Kit (Qiagen, 205313) with random priming. RNA-seq libraries were prepared with the TruSeq Stranded mRNA kit (Illumina, 20020595) and sequenced on an Illumina platform as 100 bp paired-end reads at a depth of 25 million reads/sample.

### iPSC Editing and Culture

The use of human iPS cells was approved by the McGill University Research Ethics Board (institutional review board study number A03-M19-22A). The male control/WT line used is AIW002-02 (CBIGi001-A: hPSCreg ID), a line extensively characterized and used across multiple projects in the group ^40, 41, 61, 62, 63, 64^. Cells were maintained in a 37 °C incubator with 5% CO2.

To edit the USP15 gene in the iPSC line AIW002-02, a CRISPR-Cas9 gene editing approach was employed as previously described ^41^. To generate USP15 mutant, iPSCs were nucleofected with ribonucleoprotein complex containing Cas9 protein, single guide RNA (sequence: AGAAAGAUCUUUUCUUGCUU) and homology DNA repair template (sequence: AAATTTTAAAGAAAAAATATGTTTTCCACCTTAGAAAGATCTTTTCGAGCTTTAGATTGGGATCCTGATTTGAAAAAAAGATATTTTGATGAAAATGCT) using a Lonza 4D-Nucleofector device. To generate USP15 KO cells, two gRNAs (CCAGAGUUAUCAAUGGGUCC and CACAUUUACAACUUGACUGA) targeted exon 2 and in intron 2 were used in ribonucleofection, yielding a frameshift deletion. Following limiting dilution, gene-edited clones were identified by droplet digital PCR (QX200™ Droplet Reader, Bio-Rad) and Sanger sequencing.

### Microglial Differentiation

iPSC-derived microglia were generated using a staged differentiation protocol as previously described^43^. Briefly, iPSCs were differentiated into hematopoietic progenitor cells (iHPCs) using the STEMdiff™ Hematopoietic kit (STEMCELL Technologies). On day 10, floating iHPCs were collected, centrifuged (300 g, 5 min), and either cryopreserved (Bambanker, Fujifilm Wako Chemicals) or used directly for microglial differentiation.

For microglial differentiation, iHPCs were resuspended at 5–10 × 10⁵ cells/mL in differentiation medium (MEMα supplemented with GlutaMAX, B27, Insulin-Transferrin-Selenium, Penicillin/Streptomycin, IL-34 [100 ng/mL], TGF-β1 [50 ng/mL], M-CSF [25 ng/mL], and GM-CSF [5 ng/mL]; all Thermo Fisher Scientific or Peprotech) and plated on Matrigel™-coated 6-well plates as previously described^43^. CX3CL1 (100 ng/mL) was added at day 25, and cells were considered mature at day 28. Cells were used for downstream experiments between days 28 and 42 and detached using PBS with 2 mM EDTA (10-min incubation) when needed. Differentiated microglia were validated by immunostaining for Iba1, PU.1, and TREM2.

### Generation of Human iPSC-Derived Astrocytes

Human Astrocytes were derived from iPSCs as previously described ^44^. Briefly, iPSCs at low passage were cultured on Matrigel-coated dishes in mTeSR medium until 70–80% confluence (DIV0). Cells were dissociated using Gentle Dissociation Reagent (STEMCELL Technologies), centrifuged, and replated onto Matrigel-coated 6-well plates in mTeSR medium supplemented with the ROCK inhibitor Y-27632. On DIV1, medium was replaced with Neural Induction Medium 1 [DMEM/F12 with N2, B27, BSA, SB431542, Noggin, GlutaMax, non-essential amino acids (NEAA), laminin, penicillin, and streptomycin] and refreshed every two days until DIV7. At DIV7, Neural Induction Medium 2 (same as above without SB431542 or Noggin) was applied and changed daily until DIV12, when cells were dissociated with Accutase (STEMCELL Technologies) and transferred to low-attachment dishes in Neural Progenitor Cell (NPC) Expansion Medium [DMEM/F12 with N2, B27, EGF, FGF2, GlutaMax, NEAA, laminin, penicillin, and streptomycin] with Y-27632. Neurospheres that formed within 2–3 days were harvested using a 40 μm strainer and replated onto Matrigel-coated dishes for further expansion. NPCs were stored in liquid nitrogen for later use. For astrocyte differentiation, NPCs were seeded at 15,000 cells/cm² in T25 flasks in NPC Expansion Medium containing Y-27632, and the next day switched to Astrocyte Differentiation Medium 1 (ScienCell Astrocyte Growth Medium supplemented with astrocyte growth supplement, 1% fetal bovine serum, penicillin, and streptomycin). Half of the medium was replaced every 3–4 days for 30 days, without passaging. At day 30, proliferative astrocytes were either cryopreserved in liquid nitrogen or further matured in Astrocyte Differentiation Medium 2 (FBS-free but with astrocyte growth supplement) for an additional 1–2 months, after which cells were monitored for GFAP, S100β, and aquaporin-4 expression.

### RNA isolation and real-time quantitative PCR

Total cellular RNA was isolated using the RNeasy96 kit according to the manufacturer’s protocol. RNA concentration was determined by spectrophotometry using a Nanodrop instrument (Thermo Fisher Scientific, ND-1000). Equal amounts of RNA were reverse transcribed using the cDNA reverse transcription kit (M-MLV RT Kit, Invitrogen) according to the manufacturer’s instructions on a thermocycler using the following incubation steps: 10 min at 25 °C, 1 h at 37 °C, and 7 min at 95 °C. Real-time quantitative PCR (RT-qPCR) was performed on diluted cDNA using PowerUp SYBR Green Supermix (Biorad). Gene expression was quantified using human-specific primers for MX1 (F: GGCTGTTTACCAGACTCCGACA; R: CACAAAGCCTGGCAGCTCTCTA), MX2 (F: AAAAGCAGCCCTGTGAGGCATG; R: GTGATCTCCAGGCTGATGAGCT), IRF7 (F: CCACGCTATACCATCTACCTGG; R: GCTGCTATCCAGGGAAGACACA), IFIT3 (F: CAGACAGGAAGACTTCTGAAGAAC; R: GCATTTCAGCTGTGGAAGGATT), USP18 (F: TGGACAGACCTGCTGCCTTAAC; R: CTGTCCTGCATCTTCTCCAGCA), and TBP (F: TGTATCCACAGTGAATCTTGGTTG; R: GGTTCGTGGCTCTCTTATCCTC) as the housekeeping gene. Differential expression was calculated using the 2−ΔΔCT method, with the selected housekeepers as normalizing controls. The baseline condition was defined in each experiment as RNA extracted from untreated control. All reactions were performed in technical triplicate.

### Parkinson’s disease GWAS summary statistics

The primary PD GWAS was the GP2 European 2025 release ^36^. We used the broad-European meta-analysis (file GP2_ALL_EUR_ALL_DATASET_HG38_12162024.txt.gz; 20 cohorts comprising European non-Finnish, Finnish and Ashkenazi Jewish populations across both biobank and case-control study designs), filtered as released to MAF > 0.01, *I²* < 80% and presence in ≥50% of contributing studies. Variants were aligned to GRCh38. For sensitivity analysis we additionally used the EUR-only 16-cohort subset (GP2_EUR_ONLY) and, where required, the multi-ancestry meta-analysis of Kim *et al.* ^18^. Earlier published PD GWAS leads were retrieved from Nalls *et al.* ^19^ for cross-referencing.

### Cis-eQTL summary statistics

Primary cis-eQTL data were drawn from the eQTL Catalogue ^65^. For monocytes we used the Quach *et al.* ^47^ study, with all five available 6 h stimulation panels (naive [QTD000409], LPS [QTD000414], Pam3CSK4 [QTD000419], R848 [QTD000424] and influenza [QTD000429]) at the chr12:62 Mb region. Brain and broader tissue eQTLs were drawn from GTEx v8 ^66^ and from MetaBrain ^67^. For whole-blood replication, we additionally used eQTLGen ^68^. Replication monocyte cohorts were BLUEPRINT ^48^, DICE ^49^ and OneK1K ^69^ where available within the eQTL Catalogue.

### Linkage disequilibrium reference panel and phased haplotypes

LD and phased two-locus haplotype frequencies were estimated from UK Biobank ^70^ imputed genotypes restricted to unrelated individuals of European ancestry (*n* = 339,334 individuals; *n* = 678,668 chromosomes). We extracted the chr12:61.0–63.5 Mb interval (*USP15* ±1 Mb, GRCh38) and computed pairwise *r²* and signed *r* from phased haplotype counts using plink2 ^71^. Five-marker pairwise LD and two-locus chromosome-frequency tables were also computed directly from phased imputed haplotype dosages using pgenlib (without model-based phase imputation) for the haplotype analysis presented in Supplementary Note 1.

### Conditional and joint analysis

Independent association signals at the locus were identified using GCTA-COJO ^72^. The cojo-slct procedure was run with the GP2 European 2025 summary statistics over chr12:61.0–63.5 Mb and the matched UKB LD reference, at three significance thresholds (5×10⁻⁸, 1×10⁻⁶ and 1×10⁻⁵) to assess robustness of the independent-signal nomination. Conditional analyses on each lead variant were performed with cojo-cond.

### Statistical fine-mapping

Single-locus Bayesian fine-mapping used the Wakefield approximate Bayes factor ^73^ to compute per-variant posterior inclusion probabilities (PIPs) and 95% credible sets. To accommodate multiple causal variants we additionally ran SuSiE-RSS ^74^ using the GP2 European 2025 summary statistics and the UKB LD matrix described above, with up to *L* = 5 causal effects.

### Colocalization analysis

Colocalization between PD GWAS and each candidate gene’s cis-eQTL signal was assessed with coloc.abf ^75^ and, for multi-causal cis-eQTL signals, with coloc.susie ^76^. Standard prior probabilities (*p*₁ = 1×10⁻⁴, *p*₂ = 1×10⁻⁴, *p*₁₂ = 1×10⁻⁵) were used unless otherwise stated. Robustness of PP.H4 to the choice of p₁₂ was assessed across 1×10⁻⁷ to 1×10⁻⁴ (Supplementary Note 1). We additionally ran conditional colocalization, splitting the PD GWAS by each COJO-lead via cojo-cond and re-running coloc.abf against each independent signal to test for shared causal variants between subsignals. The same coloc procedure was applied in BLUEPRINT ^48^ and DICE ^49^ for cross-cohort replication.

### Mendelian randomization

Two-sample Mendelian randomization was performed in the TwoSampleMR framework ^77^, in every Quach 2016 monocyte panel for which *USP15* cis-eQTL summary statistics are available (naive, Pam3CSK4-stimulated, influenza-stimulated; the LPS and R848 panels do not include *USP15* cis-eQTL associations because *USP15* was filtered at the per-condition gene-level QC stage in the eQTL Catalogue). cis-eQTL instruments were selected at *P* < 5×10⁻⁸ with distance pruning ≥ 100 kb between instruments. Effect alleles were harmonized between exposure and outcome. To account for both this correlation among instruments and horizontal pleiotropy through neighbouring genes in the cis-window, primary causal inference was based on the pleiotropy-robust MR-link-2 method ^78^ at the recommended default eigenvalue truncation (variance explained = 0.99), using an LD matrix computed from 10,000 subsampled UKB unrelated European individuals. For gene-attribution sensitivity, multivariable MR with *USP15* and *TAFA2* as joint exposures ^79^ was attempted across a sweep of cis-eQTL *P* thresholds (5×10⁻⁶, 10⁻⁴, 10⁻³) and pruning thresholds (*r²* < 0.05–0.50); identification was assessed by the conditional *F*-statistic ^79^. Cross-cohort MR replication was run with the same cis-MR methodology using BLUEPRINT, DICE cis-eQTL instruments separately.

### Statistical Analysis

Statistical analyses were performed using GraphPad Prism11 software. Survival curves were analyzed using Log-rank (Mantel–Cox) tests. p < 0.05 was considered statistically significant. Group comparisons for qPCR data were performed using one-way ANOVA with appropriate post-hoc tests for multiple comparisons. A p-value < 0.05 was considered statistically significant.

## DATA AVAILABILITY

The RNA-seq datasets generated for this study have been deposited in the GEO database (https://www.ncbi.nlm.nih.gov/geo/) and assigned the accession numbers GSE338123 (USP15 conditional deletion), GSE338124 (M83 PD model) and GSE341482 (mouse microglia).

## Notes

### Competing Interest Statement

The authors have declared no competing interest.

### Summary of Updates

Supplementary figures were missing from the initial submission and have been added in this revised version.

