## Supplementary figures and images for "USP15 REGULATES NEUROINFLAMMATION AND DRIVES PATHOGENESIS IN SYNUCLEINOPATHIES"

### Supplementary Figure 1

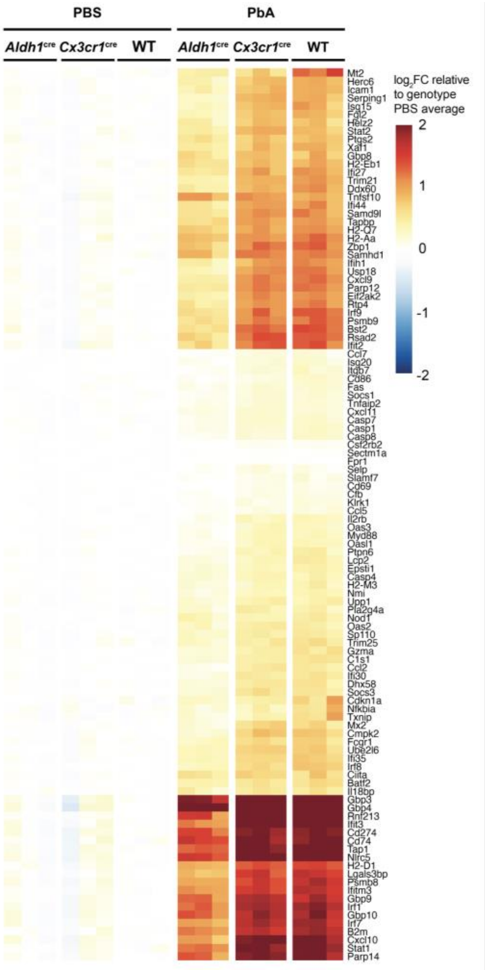


**Supplemental Figure 1**

### Supplementary Figure 2

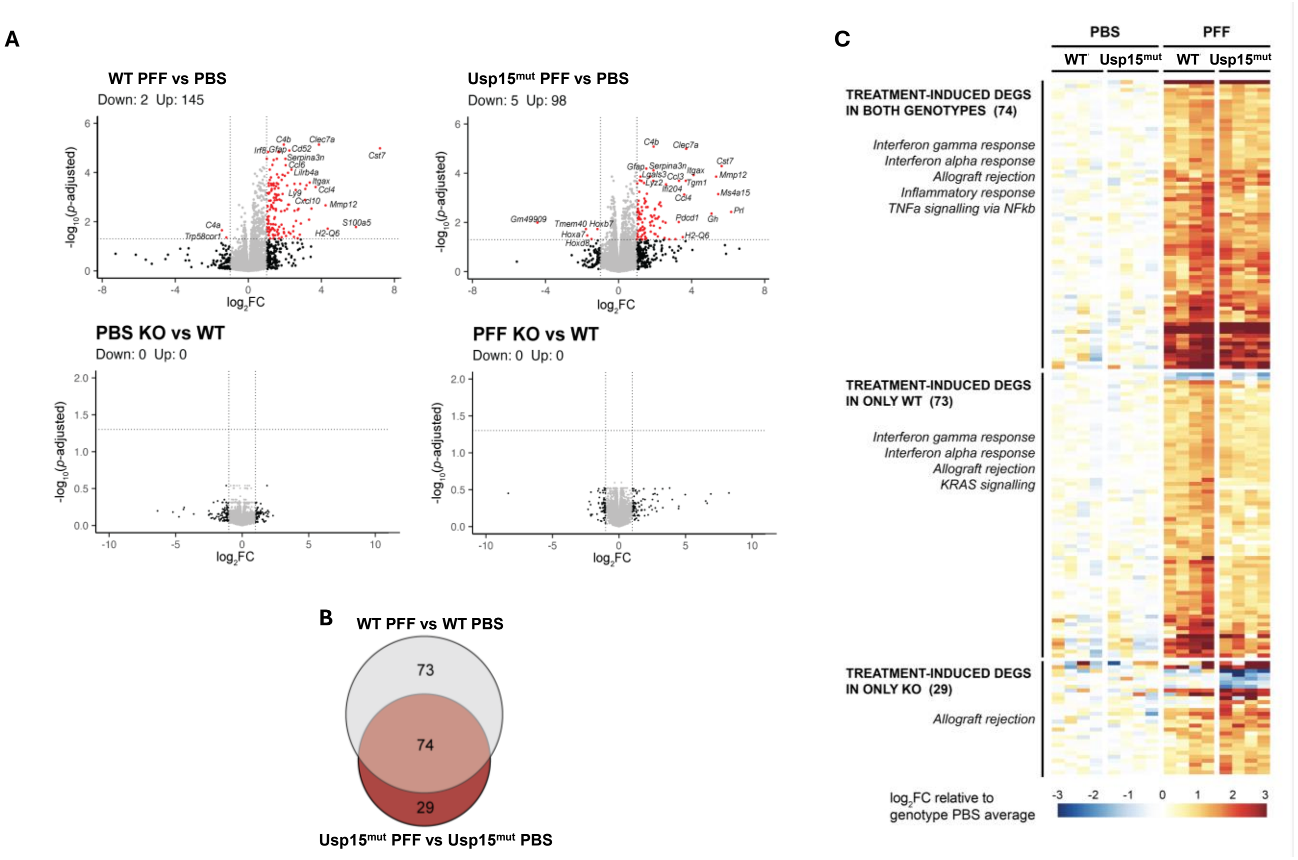


#### Supplemental Figure 2

### Supplementary Figure 3

iMGL – phagocytic activity

**A**

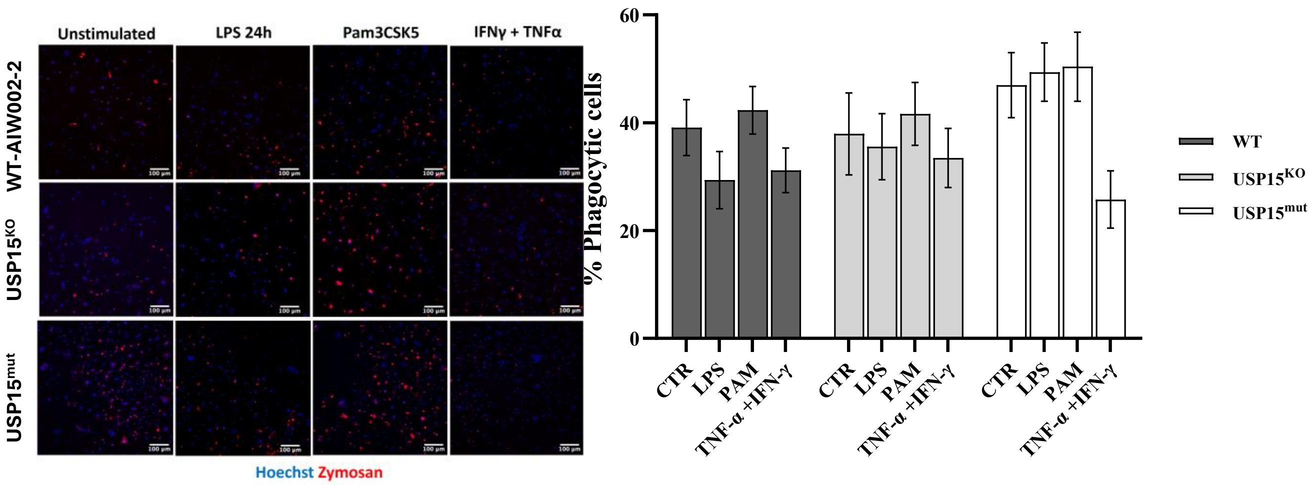


iMGL – phagocytic activity in TREM2+ cells

**B**

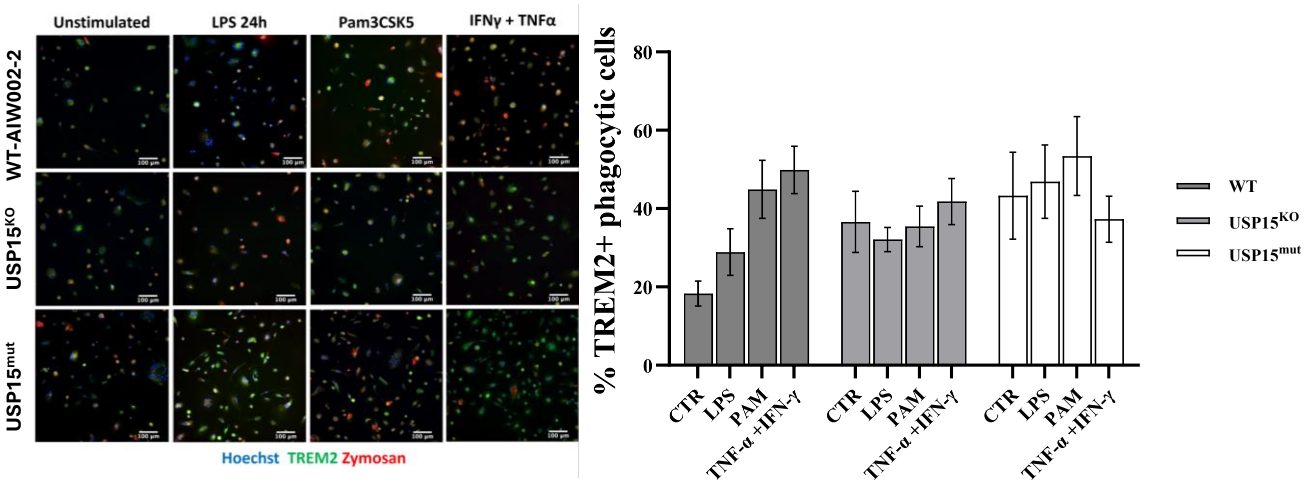


**Supplemental Figure 3**

### Supplementary Figure 4

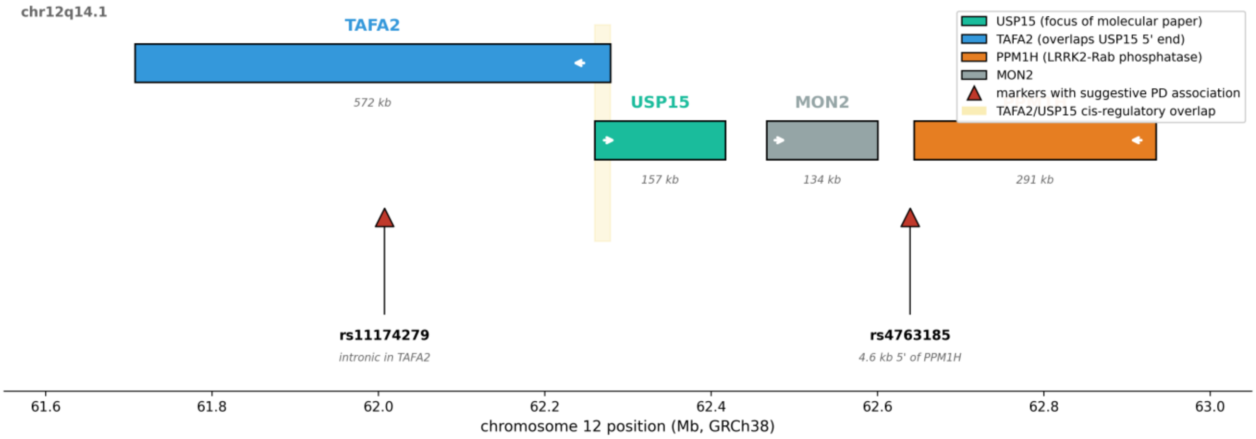


**Supplemental Figure 4**

### Supplementary Figure 5

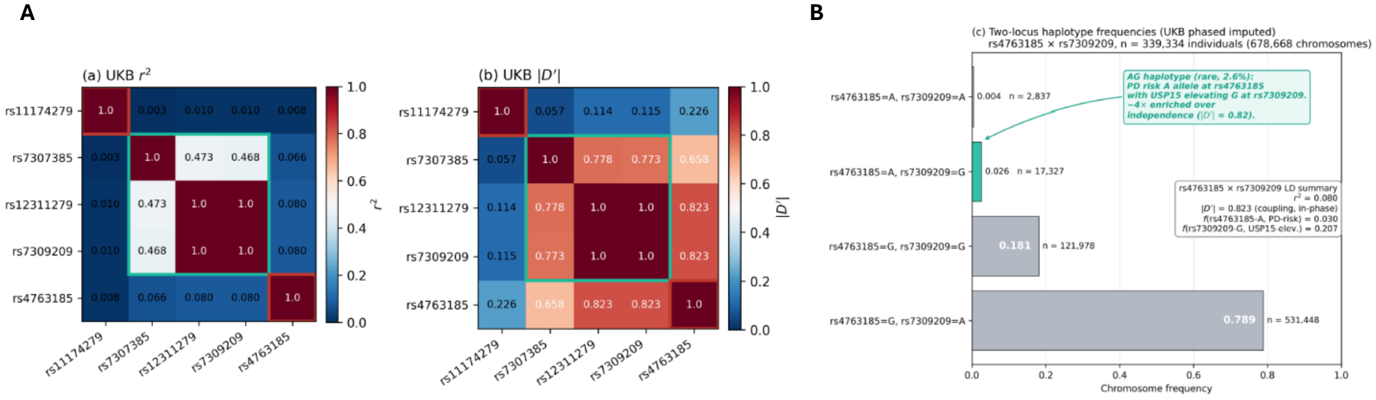


##### Supplemental Figure 5
