## Supplementary note 1 for "USP15 REGULATES NEUROINFLAMMATION AND DRIVES PATHOGENESIS IN SYNUCLEINOPATHIES"

### Genetic analyses at the chr12q14.1 USP15 locus

In the GP2 European 2025 PD meta-analysis (20 cohorts), the chr12:62 Mb interval containing *USP15*, *TAFA2*, *MON2* and *PPM1H* shows a suggestive but sub-genome-wide-significant association (lead rs4763185, $P = 3.0\times10^{-6}$, $\beta = +0.095$, EAF = 0.032). The locus is not among the genome-wide-significant loci of that study or of Nalls *et al.* 2019, but *USP15* carries a positive Polygenic Priority Score (PoPS = +0.256). GCTA-COJO on the GP2 statistics with a UK Biobank chr12 LD reference (n = 339,334 unrelated European-ancestry individuals) resolved two conditionally independent signals, rs4763185 (5′ of *PPM1H*) and rs11174279 (intronic in *TAFA2*), in essentially zero pairwise LD ($r^2 = 0.008$).

Of the four genes, only *USP15* and *TAFA2* carry monocyte cis-eQTL signals over chr12:62.1 to 62.4 Mb (Quach *et al.* 2016, n = 200), with overlapping credible sets. Colocalization with the PD GWAS gives PP.H4 ≈ 0.81 for both genes; conditioning on each COJO lead localizes the shared signal to the rs4763185 subsignal (PP.H4 = 0.71 after removing rs11174279, 0.18 after removing rs4763185), stable across priors and under an in-sample SuSiE-RSS reference. *PPM1H* and *MON2* show no coincident colocalization. The signal replicates in BLUEPRINT (n = 191; *USP15* PP.H4 = 0.80, *TAFA2* = 0.74); in the smaller DICE cohort (n = 91) the posterior is power-limited but the cis-eQTL effect sizes are direction-concordant (heterogeneity P = 0.97 at rs7309209).

In UK Biobank phased haplotypes, the rs4763185 PD-risk allele and the *USP15* cis-eQTL elevating allele at rs7309209 lie in coupling-phase LD ($|D'| = 0.82$, $r^2 = 0.08$), and the chromosome carrying the cis-eQTL allele without the rs4763185 risk allele still confers a small additive PD risk (per-chromosome OR = 1.025, 95% CI 1.007 to 1.043, $P = 5.6\times10^{-3}$), indicating a contribution independent of rs4763185 (Supplementary Figure 5).

Two-sample cis-Mendelian randomization across the Quach monocyte panels gave positive *USP15* → PD estimates, significant after pleiotropy correction (MR-link-2 $\alpha = +0.022$, $P = 6.1\times10^{-6}$ in naive monocytes; positive and significant in the Pam3CSK4 and influenza panels). Because *USP15* and *TAFA2* share most of their cis-eQTL architecture (instrument correlation $r \geq 0.98$), multivariable MR was non-identifiable (conditional $F < 2$), so the regulatory MR signal implicates the *USP15*/*TAFA2* cis-window collectively;

gene-level attribution to *USP15* rests on the molecular and mouse evidence in the main text.
