## Supplementary table 1 for "USP15 REGULATES NEUROINFLAMMATION AND DRIVES PATHOGENESIS IN SYNUCLEINOPATHIES"

### Supplementary Table 1. Gene Co-expression in PD Microglia

**USP15 / LRRK2 co-expression in PD microglia (ShinyCell)**

| Expression > 0 | nCells | % |
|---|---|---|
| Both (USP15 + LRRK2) | 1,105 | 53.33% |
| USP15 only | 638 | 30.79% |
| LRRK2 only | 156 | 7.53% |
| Neither | 173 | 8.35% |

**USP15 / SNCA (α-syn) co-expression in PD microglia (ShinyCell)**

| Expression > 0 | nCells | % |
|---|---|---|
| Both (USP15 + SNCA) | 1,119 | 54.01% |
| USP15 only | 624 | 30.12% |
| SNCA only | 159 | 7.67% |
| Neither | 170 | 8.20% |
