## Supplementary table 2 for "USP15 REGULATES NEUROINFLAMMATION AND DRIVES PATHOGENESIS IN SYNUCLEINOPATHIES"

**Supplementary Table 2. Two-locus haplotype frequencies and additive PD odds ratios (rs4763185 x rs7309209)**

| Haplotype | rs4763185 | rs7309209 | Frequency | N_chromosomes | OR | OR_95CI_low | OR_95CI_high | P |
|---|---|---|---|---|---|---|---|---|
| GA | G | A | 0.789 | 531,448 | 1.000 | | | (reference) |
| GG | G | G | 0.181 | 121,978 | 1.025 | 1.007 | 1.043 | 0.0056 |
| AA | A | A | 0.004 | 2,837 | 1.083 | 1.039 | 1.128 | 1.4E-04 |
| AG | A | G | 0.026 | 17,327 | 1.110 | 1.066 | 1.155 | 3.6E-07 |

*Two-locus haplotype counts at rs4763185 and rs7309209 were derived from UK Biobank imputed phased genotypes (release v3) on n = 339,334 unrelated European-ancestry individuals (678,668 chromosomes). Counts are tabulated over the 336,795 individuals (673,590 chromosomes) with non-missing imputed phased calls at both loci; 2,539 individuals (5,078 chromosomes; 2,261 missing at rs4763185 and 282 at rs7309209) with imputation missingness at either locus were excluded from the two-locus haplotype tabulation.*

*OR = 1.000 marks the reference (common-background) haplotype. Per-haplotype ORs from a two-SNP joint solve of the GP2 European meta-analysis marginal effects.*
