## Supplementary table 3 for "USP15 REGULATES NEUROINFLAMMATION AND DRIVES PATHOGENESIS IN SYNUCLEINOPATHIES"

### Supplementary Table 3. Cross-cohort cis-eQTL colocalization and Mendelian randomization

| Cohort | N | USP15_PP.H4 | TAFA2_PP.H4 | USP15_MR_P | TAFA2_MR_P |
|---|---|---|---|---|---|
| Quach naive | 200 | 0.810 | 0.810 | 1.9E-08 | 1.6E-08 |
| BLUEPRINT | 191 | 0.800 | 0.740 | 0.0035 | 8.6E-04 |
| DICE | 91 | 0.120 | 0.380 | n.s. | n.s. |

*PP.H4: posterior probability of a shared causal variant (coloc.abf, default priors) between the GP2 European 2025 PD GWAS and each gene's monocyte cis-eQTL.*

*MR_P: two-sample cis-Mendelian-randomization P. n.s. = not significant (DICE is power-limited, n = 91; cis-eQTL effect sizes remain direction-concordant, heterogeneity P = 0.97 at rs7309209).*
