## Supplementary table 4 for "USP15 REGULATES NEUROINFLAMMATION AND DRIVES PATHOGENESIS IN SYNUCLEINOPATHIES"

### Supplementary Table 4. cis-eQTL Mendelian randomization across Quach 2016 monocyte

| Panel | N | IVW_N_instruments | IVW_alpha | IVW_P | Bonferroni_3test_P | MRlink2_alpha | MRlink2_P |
|---|---|---|---|---|---|---|---|
| naive | 200 | 2 | 0.074 | 1.9E-08 | 5.6E-08 | 0.022 | 6.1E-06 |
| Pam3CSK4 (6h) | 196 | 0 | n/a | n/a | n/a | 0.012 | 0.0056 |
| Influenza (6h) | 198 | 2 | 0.057 | 1.5E-08 | 4.4E-08 | 0.007 | 0.02 |

*IVW: inverse-variance-weighted two-sample MR; alpha on the log-OR scale per allele. Bonferroni_3test_P corrects across the three panels.*

*MR-link-2: pleiotropy-robust estimator; pleiotropy term significant in every panel.*

*Multivariable MR (USP15 and TAFA2 as joint exposures) was non-identifiable in every configuration tested (conditional F < 2; cis-eQTL instrument effect-size correlation r >= 0.98).*

*The LPS and R848 panels are absent because USP15 cis-eQTL summary statistics were filtered at the per-condition gene-level QC stage in the eQTL Catalogue.*
