## Supplementary table 5 for "USP15 REGULATES NEUROINFLAMMATION AND DRIVES PATHOGENESIS IN SYNUCLEINOPATHIES"

**Supplementary Table 5. Three-locus haplotype frequencies and additive PD odds ratios (rs11174279 x rs4763185 x rs7309209)**

| Haplotype | rs11174279 | rs4763185 | rs7309209 | Frequency | N_chromosomes | OR | OR_95CI_low | OR_95CI_high | P |
|---|---|---|---|---|---|---|---|---|---|
| TGA | T | G | A | 0.662 | 449,470 | 1.000 | | | (reference) |
| TGG | T | G | G | 0.137 | 92,906 | 1.021 | 1.004 | 1.039 | 0.016 |
| AGG | A | G | G | 0.112 | 76,037 | 0.961 | 0.944 | 0.978 | 1.4E-05 |
| AGA | A | G | A | 0.041 | 28,052 | 0.981 | 0.956 | 1.007 | 0.155 |
| TAG | T | A | G | 0.018 | 12,101 | 1.111 | 1.068 | 1.156 | 2.0E-07 |
| AAG | A | A | G | 0.007 | 5,050 | 1.068 | 1.022 | 1.115 | 0.003 |
| AAA | A | A | A | 0.003 | 2,036 | 1.045 | 1.001 | 1.092 | 0.046 |
| TAA | T | A | A | 0.001 | 748 | 1.088 | 1.045 | 1.133 | 5.0E-05 |

*Alleles in column order rs11174279, rs4763185, rs7309209. Counts from UK Biobank phased imputed haplotypes (n = 339,334 individuals; 678,668 chromosomes).*

*Per-haplotype log-OR is the additive sum of joint allele effects from a three-SNP GCTA-COJO solve on GP2 European meta-analysis summary statistics (UK Biobank chr12 LD reference): beta(rs11174279-A) = -0.040 (P = 1.4e-5); beta(rs4763185-A) = +0.084 (P = 5.0e-5); beta(rs7309209-G) = +0.021 (P = 1.6e-2).*
